# Receptor endocytosis orchestrates the spatiotemporal bias of β-arrestin signaling

**DOI:** 10.1101/2023.04.27.538587

**Authors:** András Dávid Tóth, Bence Szalai, Orsolya Tünde Kovács, Dániel Garger, Susanne Prokop, András Balla, Asuka Inoue, Péter Várnai, Gábor Turu, László Hunyady

**Affiliations:** Institute of Enzymology, Centre of Excellence of the Hungarian Academy of Sciences, Research Centre for Natural Sciences, Eötvös Loránd Research Network, Budapest, Hungary; Department of Physiology, Faculty of Medicine, Semmelweis University, Budapest, Hungary; Department of Internal Medicine and Haematology, Semmelweis University, Budapest, Hungary; Computational Health Center, Helmholtz Munich, Neuherberg, Germany; ELKH-SE Laboratory of Molecular Physiology, Eötvös Loránd Research Network, Budapest, Hungary; Molecular and Cellular Biochemistry, Graduate School of Pharmaceutical Sciences, Tohoku University, Sendai, Japan

## Abstract

The varying efficacy of biased and balanced agonists is generally explained by the stabilization of different active receptor conformations. In this study, systematic profiling of transducer activation of AT_1_ angiotensin receptor agonists revealed that the extent and kinetics of β-arrestin binding exhibit substantial ligand-dependent differences, which however completely disappear upon the inhibition of receptor internalization. Even weak partial agonists for the β- arrestin pathway acted as full or near full agonists, if receptor endocytosis was prevented, indicating that receptor conformation is not an exclusive determinant of β-arrestin recruitment. The ligand-dependent variance in β-arrestin translocation at endosomes was much larger than it was at the plasma membrane, showing that ligand efficacy in the β-arrestin pathway is spatiotemporally determined. Experimental investigations and mathematical modeling demonstrated how multiple factors concurrently shape the effects of agonists on endosomal receptor–β-arrestin binding and thus determine the extent of bias. Among others, ligand dissociation rate and G protein activity have particularly strong impact on receptor–β-arrestin interaction, and their effects are integrated at endosomes. Our results highlight that endocytosis forms a key spatiotemporal platform for biased GPCR signaling and can aid the development of more efficacious functionally-selective compounds.

****One Sentence summary**:** Agonist-specific differences in β-arrestin recruitment are mainly determined by the ligand dissociation rate and G protein activation at the endosomes.

## INTRODUCTION

G protein-coupled receptors (GPCRs) represent the largest family of cell surface receptors and engage a variety of signaling proteins upon agonist stimulation. Certain ligands are known to induce selective or stronger activation of different transducers, a phenomenon called biased signaling (*1*). The concept has gained great attention as biased drugs may exert beneficial clinical effects, because they may not engage signaling pathways that induce undesired side effects. Regarding biased signaling, the AT_1_ angiotensin receptor (AT_1_R) is one of the most extensively studied receptors. Whereas its endogenous peptide ligand angiotensin II (AngII) serves as a full agonist for AT_1_R, several studies have shown that derivatives of AngII, which lack an aromatic amino residue in the 8^th^ position, prefer β-arrestin over G protein activation (*2, 3*). Moreover, diverse functional actions of AT_1_R agonists have also been demonstrated across different G protein and G protein-coupled receptor kinase (GRK) subtypes (*4–7*). Above all, different ligand bias profiles have also been linked to specific *in vivo* effects and TRV120027, a β-arrestin–biased agonist, has been even evaluated in clinical trials (*8–12*).

It was theorized that the pathway-selective cellular actions of biased ligands are based on their ability to stabilize receptors in different conformations (*13, 14*), which was later proven by the elucidation of the corresponding crystal structures (*15*). In line with a recent guideline, here we use the term “ligand bias” to refer to biased signaling emerging from distinct agonist-induced receptor conformations (*1*). Despite the unique translational potential, it has remained elusive how ligand bias interferes with the generally known kinetic and spatial factors that regulate receptor signaling. Recent advancements in live cell-based sensors and genetically modified cell lines have greatly improved our understanding of how the temporal alteration or synchronization of signaling pathways can transmit specific information (*16, 17*). Moreover, the concept and importance of “temporal bias” is increasingly acknowledged, as many studies have pointed out that the activity of distinct signaling pathways can differentially evolve over time, and the kinetics of these changes happen in a ligand-specific manner (*1, 17–19*). Besides, data are emerging that some ligands exhibit location or spatial bias, which means that they may differently induce receptor signaling in distinct subcellular compartments (*20–23*). These levels of complexity pose a great challenge to the precise experimental investigation of the kinetic and spatial factors that impact biased signaling, and consequently complicate the rational design of novel pathway-selective clinical drugs.

In this study, we aimed to identify the principal dynamic processes that act interdependently with ligand bias to evoke functionally selective cellular responses. For the comprehensive investigation of the spatiotemporal layer of biased signaling, we conducted a systematic series of advanced kinetic assays with a set of AT_1_R agonists and formulated an in silico model of receptor signaling. We found that differences in the extent of AT_1_R–β-arrestin interaction upon distinct agonist stimuli, including balanced and biased ligands, almost completely disappear upon inhibition of receptor internalization, and the key regulatory factors that drive ligand specificity in β-arrestin binding are integrated at endosomes. Our results reveal a strong correlation between the ligand dissociation rate and the extent of AT_1_R–β-arrestin interaction after receptor internalization. Furthermore, our data reveal that the β-arrestin2 recruitment of balanced agonists is enhanced by G_q/11_ activity, predominantly at the endosomal compartment. Finally, our mathematical model and expanded experimental results with β_2_-adrenergic receptor imply that endocytosis provides a general platform for the kinetic and spatial factors to shape the overall signaling outcome together with ligand bias, in a mutually dependent manner.

## RESULTS

### Ligand-specific differences in AT_1_R–β-arrestin binding depend on receptor endocytosis

To take a comprehensive look at the temporal characteristics of biased signaling, we real-time monitored the activation of a large set of AT_1_R transducers after stimulation with 9 AT_1_R peptide ligands, which are known to display markedly different affinities and signaling bias profiles **(Fig. S1)** (*24–26*). All agonists were applied at 10 µM concentration, which results in complete or near complete AT_1_R saturations, and the endogenous agonist angiotensin II (AngII) was selected as the reference ligand. In bioluminescence resonance energy transfer (BRET) measurements between RLuc-labeled AT_1_R and Venus-tagged β-arrestins, all ligands were able to induce β-arrestin1 (βarr1, **Fig. S2**) and β-arrestin2 (βarr2, **Fig. 1A**) binding to AT_1_R, however their efficacies varied. We found that the differences in agonist effects continuously increased over time, i.e. fold difference between AngII and SII-AngII was 1.6-fold at 2 min, while it was 3.2-fold at 20 min after stimulation for β-arrestin2 recruitment (**Fig. 1A**). In contrast to β-arrestin recruitment, the ligands could be divided into two groups based on their ability to activate the G_q_ protein, evaluated using the TRUPATH BRET biosensor (**Fig. 1B**) (*27*). These groups are referred to as G_q_-activating and non-G_q_-activating ligands. The latter group is also frequently named as β-arrestin-biased agonists. In addition, G_q_-activating peptides effectively activated other G protein TRUPATH sensors (G_11_, G_i1_, G_i2_, G_i3_, G_oA_, G_oB_, G_12_, and G_13_) as well, however, some G proteins were also partially activated by the β-arrestin-biased agonists (**Fig. S3**). The activation kinetics of the distinct G protein sensors greatly differed, however, in contrast to the diverging ligand-dependent kinetics of β-arrestin binding, the relative differences between ligand effects were stable over time (**Fig. 1B**–**C**). These data are consistent with previous observations that AT_1_R can be stabilized in multiple active conformations, which may selectively couple to distinct transducers. In addition, our results demonstrate the existence of profound temporal differences, which influence signaling efficacy in a ligand-and transducer-specific manner, and, thus, shape the extent of the observed bias.

**Figure 1.**
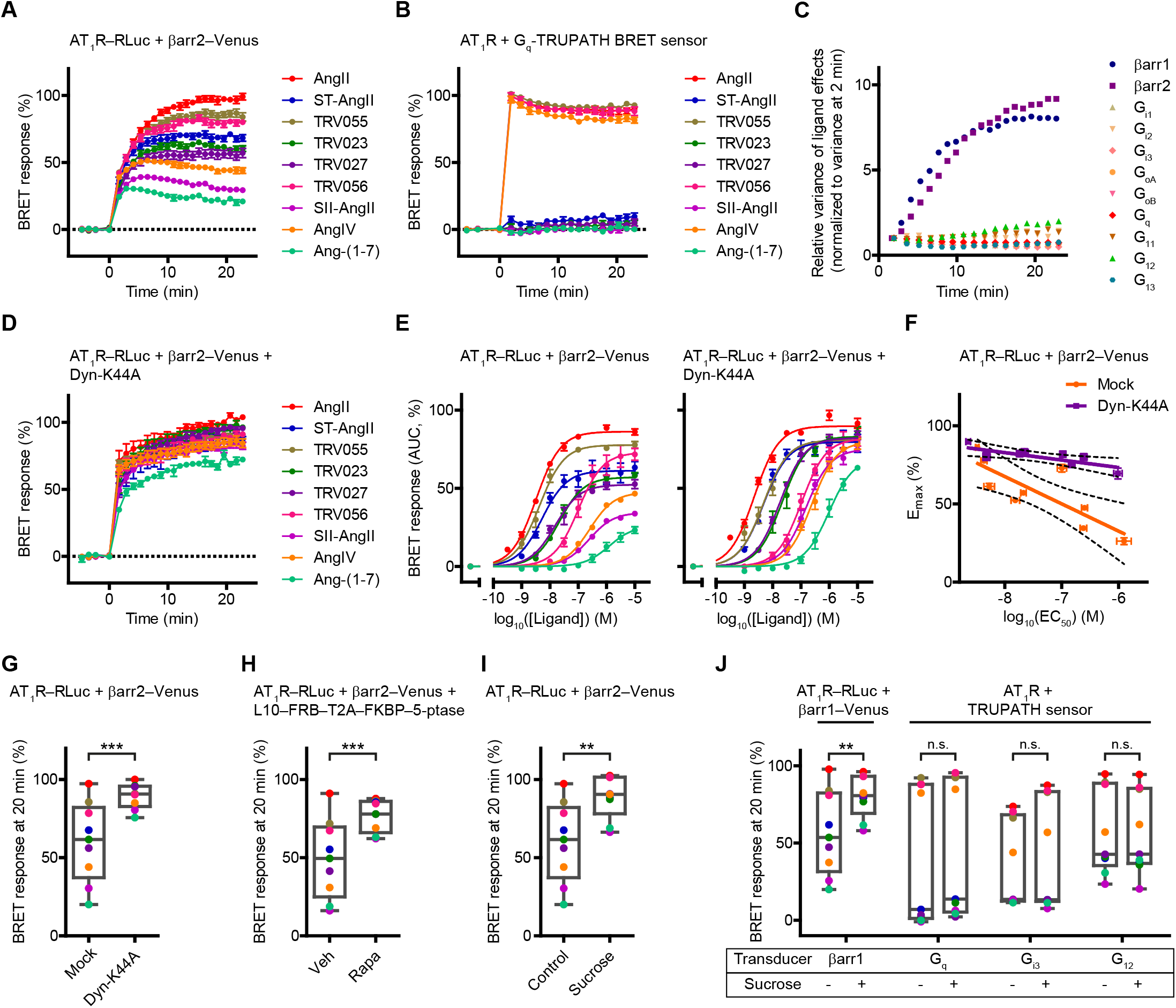
Ligand-specific temporal differences in AT_1_R–β-arrestin binding disappear upon the inhibition of receptor internalization. **A** Real-time measurement of AT_1_R–β-arrestin2 binding upon agonist stimulation. The agonists showed varying effectiveness, and the ligand-specific differences increased over time. **B** Real-time measurement of G_q_ protein activation after AT_1_R stimulation. In contrast to β-arrestin recruitment, agonists formed two discrete groups based on their ability to activate the G_q_ biosensor. The same color code is used as in A. **C** Analysis of the variance of ligand-specific effects over time. The variance of ligand-specific BRET responses (squared standard deviation of BRET responses of all agonists) was calculated and were normalized to the variance detected at the first measurement point (∼2 min) after stimuli in order to display the temporal change in the distribution of ligand-specific signals. In the case of G protein activation, the differences between agonists already appeared at the first time point and remained almost the same during the whole investigation period. In contrast, the ligand-dependent differences gradually rose in the case of β-arrestin recruitment. The corresponding kinetic curves are shown in Fig. 1A–B, Fig. S2A, Fig. S3A–H. **D** Effect of Dyn-K44A overexpression on AT_1_R–β-arrestin2 interaction. In contrast to the control condition (A), the divergence of the binding curves over time markedly decreased. Dyn-K44A overexpression abolished the agonist-specific differences in β-arrestin2 binding efficacy and kinetics. **E** Concentration–response curves of AT_1_R–β-arrestin2 binding with or without Dyn-K44A co-expression, area under curve (AUC) values are shown. The corresponding kinetic curves are shown in Fig. S4 and S5, and the fitted EC_50_ values are shown in Fig. S5J. **F** Relationship between the potency and efficacy values of different agonists in the presence or absence of receptor internalization. Data display linear correlation under both conditions (Mock: r^2^=0.6624; Dyn-K44A: r^2^=0.5685), but the slope of the linear regression curves was significantly different (−17.45±4.708 vs. −4.745±1.506, *, *P* = 0.0233). 95% confidence intervals are marked by dotted lines. **G**–**I** Inhibition of endocytosis by three independent experimental methods confirm the significant decrease of ligand-dependent differences in β-arrestin binding at later (∼20 min) measurement time points: overexpression of Dyn-K44A (G) ***, P = 0.0002; PtdIns(4,5)P_2_ depletion with rapamycin (H) ***, P = 0.0009; or hypertonic sucrose treatment (I) **, P = 0.0083, F-tests were performed to evaluate statistical significance, data were normalized to the averages in order to statistically compare the standard deviations of each distribution. Box and whiskers with aligned dot plots show distribution of the mean response magnitudes. **J** The binding pattern of β-arrestin1 is similarly affected by the blockade of endocytosis as that of β-arrestin2. However, the activity profiles of different ligands regarding G protein activation are unchanged. Data are processed and displayed as in G–I. **, P = 0.0077 for βarr1, P = 0.7385 for G_q_, P = 0.7993 for G_i3_, P = 0.9681 for G_12_.The corresponding kinetic curves are shown in Fig. 1A and 1D (for G); Fig. S6B (for H), Fig. 1A and S7A (for I), Fig. S2A and S7C; 1B and S7D; S3D and S7D; S3G and S7F (for J). Except for the concentration–response curves, the ligands were applied at 10 μM. *N* = 3–19. Data are mean ± SEM in A–F. Data were expressed as a percentage of the peak AngII (100 nM or 10 μM)-induced effect of the kinetic curves in the same expression condition (i.e. with or without Dyn-K44A coexpression). In addition, since sucrose altered the luminescence intensities, the changes in BRET ratio of sucrose pretreated cells were expressed as a percentage of AngII-induced effect after sucrose pretreatment.

Since a substantial pool of receptors are expected to be internalized in the investigated time frame, we assessed how their spatial distribution influences the temporal aspects of transducer activity. First, we focused on the β-arrestin pathway, where the most prominent temporal differences were observed. To study this question, we overexpressed a dominant negative form of dynamin2A (Dyn-K44A) to inhibit endocytosis (*28*). Remarkably, we observed that the ligand-dependent differences in AT_1_R–β-arrestin2 binding almost completely disappeared in Dyn-K44A expressing cells (**Fig. 1D** vs. **Fig. 1A**). For instance, the prototypical β-arrestin-biased agonist SII-AngII turned from a weak partial agonist to a near full agonist. Concentration–response analysis revealed a strong relation between the efficacy and potency values of distinct agonists under normal conditions. However, when endocytosis was inhibited, we found almost equal ligand efficacies in β-arrestin2 recruitment (**Fig. 1E**–**G** and **S4**–**5**), whereas the potency values of distinct ligands were not significantly different (**Fig. 1F** and **S5J**).

To verify the robust effects of receptor endocytosis, we applied two additional methodologies for the inhibition of internalization. First, we used a rapamycin-inducible phosphatidylinositol-4,5-bisphosphate (PtdIns(4,5)P_2_) depletion system (**Fig. S6**), since acute degradation of plasma membrane PtdIns(4,5)P_2_ was shown to prevent GPCR internalization (*25, 29*). We found that PtdIns(4,5)P_2_-depletion did not alter the AngII-induced AT_1_R–β-arrestin2 binding, but markedly enhanced the effects of the less efficacious agonists (**Fig. S6** and **1H**). We also used hypertonic sucrose solution, a known inhibitor of clathrin-mediated endocytosis (*25, 28, 30*). Upon pretreatment with hypertonic sucrose, highly similar effects were observed on β- arrestin2 recruitment (**Fig. 1I** and **S7A).** During the experiments, we found that hypertonic sucrose per se decreased the detected luminescence intensities probably by affecting the luciferase activity (**Fig. S7B**). Nevertheless, the administration of hypertonic sucrose had the advantage that no additional construct had to be expressed, making it easy to use in different assays of transducer activation. The inhibition of receptor endocytosis with hypertonic sucrose exerted similar changes in β-arrestin1 recruitment as that of β-arrestin2 (**Fig. 1J** and **S7C**). In contrast, the activation kinetics of G_q_, G_i3_, and G_12_, representative members of G protein subfamilies, were only slightly or moderately affected, and the overall differences between ligands in their ability to activate G proteins were not altered significantly (**Fig. 1J** and **S7D**–**F**).

The effect of internalization inhibition was further tested at the second messenger level. PtdIns(4,5)P_2_ cleavage, a hallmark of G_q/11_ protein–phospholipase Cβ (PLCβ) activation, was monitored upon AT_1_R stimulation with or without Dyn-K44A overexpression. In agreement with the unchanged G_q_ biosensor activation upon hypertonic sucrose treatment, Dyn-K44A overexpression did not alter the relative effects of ligands on PtdIns(4,5)P_2_ levels (**Fig. S8**). Taken together, we concluded that receptor endocytosis determines the ligand-dependent differences in β-arrestin binding but it does not alter the inherent ability of the active receptor conformation to induce G protein activity. Thus, non-G_q_-activating ligands preserve their β-arrestin-biased property under endocytosis-inhibited conditions, but their partial agonism in β-arrestin binding turns into full or near full agonism.

### Ligand-dependent differences in AT_1_R–β-arrestin2 binding are primarily caused by the diverse ability of ligands to stabilize endosomal AT_1_R–β-arrestin2 complexes

In the next set of experiments, we aimed to elucidate how endocytosis induces ligand-dependent differences in β-arrestin binding. We hypothesized that the variability of agonist efficacies in β- arrestin recruitment is the consequence of differences in the formation of endosomal agonist– receptor–β-arrestin complexes. To study this hypothesis, we created cell compartment-targeted biosensors and monitored the β-arrestin recruitment in different compartments. We fused the BRET donor enzyme NanoLuc either to a myristoylation-palmitoylation sequence or to Rab5 in order to target it to the plasma membrane or to early endosomes (PM–NanoLuc and EE–NanoLuc) and applied these biosensors together with Venus-tagged β-arrestin2 in bystander BRET measurements (**Fig. 2A**). After AngII treatment, the BRET signal between PM–NanoLuc and β- arrestin2–Venus first increased then slightly decreased (**Fig. 2B**), which reflects the plasma membrane translocation and the concomitant trafficking of β-arrestin2–Venus. Consistent with this, we measured a slightly delayed increase in the BRET signal between EE–NanoLuc and β- arrestin2–Venus, which represents the enrichment of β-arrestin2–Venus at endosomes (**Fig. 2C**). Similar to AngII, all other AT_1_R ligands were able to induce plasmalemmal and endosomal β- arrestin2 translocations as well (**Fig. 2B**–**C**). However, the ligand-dependent differences in plasmalemmal β-arrestin2 translocation were significantly smaller compared to the differences in endosomal β-arrestin2 recruitment, and the latter was mainly responsible for the overall variance of the total AT_1_R–β-arrestin2 binding (**Fig. 2D**–**E**).

**Figure 2.**
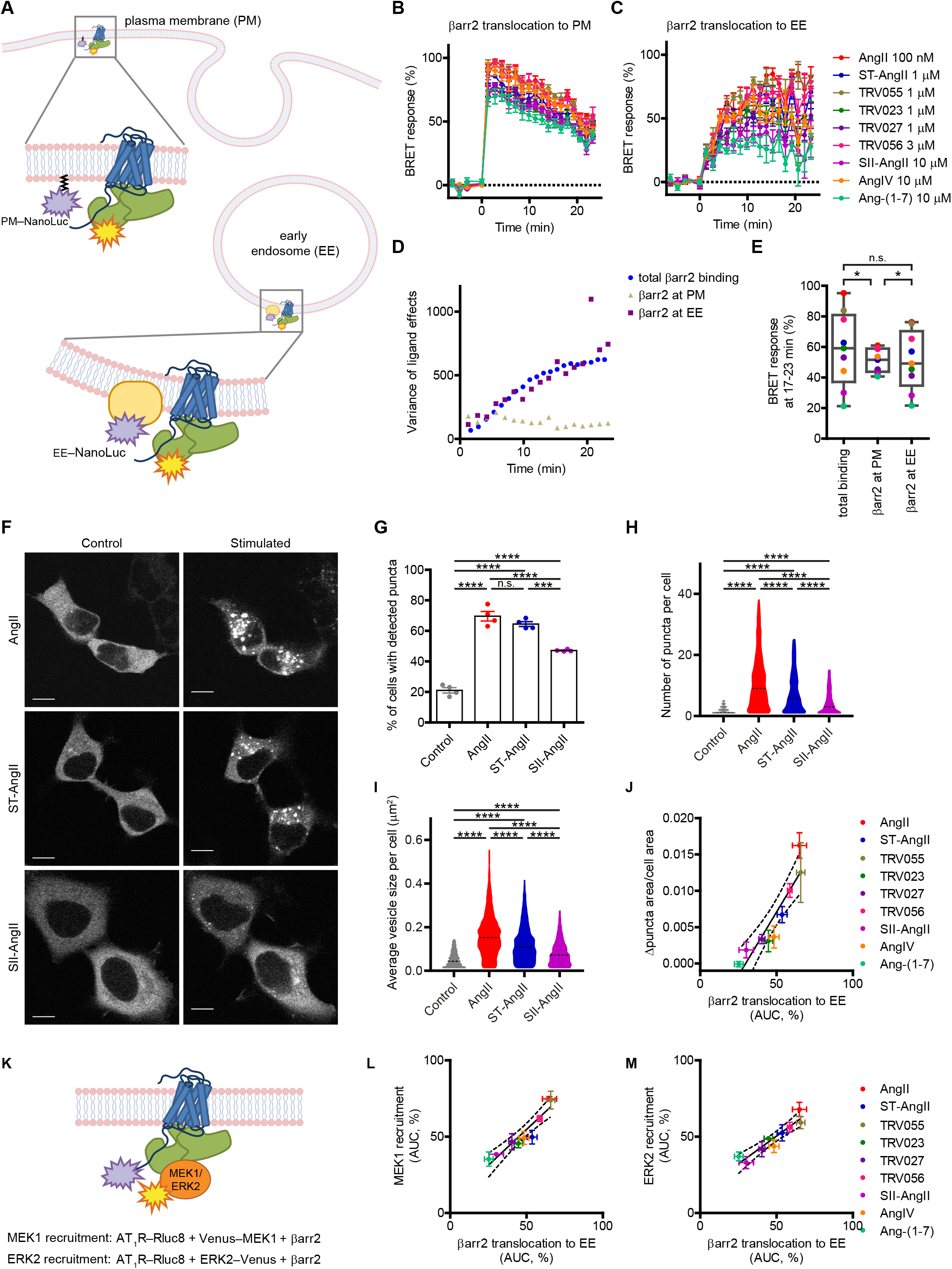
AT_1_R agonists substantially differ in their ability to induce β-arrestin2 recruitment to endosomes. **A** Schematic illustration of the BRET setups for the compartment-specific monitoring of β- arrestin2 translocation. **B–C** Real-time monitoring of β-arrestin2 translocation to the plasma membrane (PM) and to early endosomes (EEs). The indicated saturating ligand concentrations were applied, *N* = 4. **D–E** Comparison of AT_1_R–β-arrestin2 binding at different subcellular compartments. Total β- arrestin2 binding represents the BRET signal between AT_1_R–RLuc and β-arrestin2–Venus (Fig.1A). The time-dependent variability between the ligands’ effectiveness in total β-arrestin2 recruitment was not recaptured in the plasma membrane. However, a similar temporal profile of the variance (squared standard deviation of BRET responses of all agonists) was found at the endosomal compartment implicating its dominant contribution to the observed total variance. In **E**, agonist-induced β-arrestin2 responses at later time points (average of BRET responses collected 17–23 min after stimulation) are shown. The ligand-dependent variance was significantly lower at the plasma membrane than at the endosomes or that of the total binding (β-arr2 at PM vs. β-arr2 at EE: *, P = 0.0405; total binding vs. β-arr2 at PM: *, P = 0.0493; total binding vs. β-arr2 at EE: P = 0.9261), F tests were performed on average normalized datasets. **F**–**J** Endosomal β-arrestin2 translocation analyzed by confocal fluorescence microscopy. **F** Representative images of live cells expressing β-arrestin2–Venus, before and after stimulation with 10 μM AngII, ST-AngII or SII-AngII for 20–30 min. Scale bars are 10 μm. **G** Quantitative analysis of agonist-induced vesicle formation in live cells. Intracellular β- arrestin2–Venus puncta were identified by a machine learning-based algorithm, details are discussed in Methods and Fig. S9. The percentage of cells that contained fluorescent puncta are plotted. All ligands induced significant response (vs. control, ****, P < 0.0001) but with varying efficacy (AngII vs. SII-AngII, ****, P < 0.0001; ST-AngII vs. SII-AngII, ***, P = 0.0002; AngII vs. ST-AngII, P = 0.5284), one-way ANOVA with Bonferroni post-hoc test was applied. *N* = 4. **H**–**I**, Violin plots display the distribution of the number (H) and average size (I) of intracellular β- arrestin2–Venus puncta per cell. Only cells with detected puncta from 4 independent experiments were included into the analysis, 1508, 1584, 1268 and 1040 cells for Control, AngII, ST-AngII, and SII-AngII were analyzed. Outliers were identified and removed by the ROUT method (Q = 1%). The amount and the size of the β-arrestin2–Venus containing vesicles were both significantly different between the treatments (****, P <0.0001), one-way ANOVA with Bonferroni post-hoc test was used for statistical evaluation. **J** The ligand-specific extent of endosomal β-arrestin2 recruitment measured by BRET technique (C) strongly correlated with the observations of the confocal microscopic experiments. Image acquisition was performed on PFA-fixed cells, which were stimulated with the same ligand concentrations as in C, *N* = 4. The ratio of agonist-induced intracellular vesicle area and whole cell area was assessed for all AT_1_R agonists and normalized to the ratio of unstimulated cells. These values are plotted against the AUC values from the kinetic curves in panel C. Linear regression, r^2^ = 0.8492, ***, P = 0.0004, dotted lines indicate 95% confidence intervals. **K**–**M** Monitoring of the AT_1_R–β-arrestin2–MEK/ERK2 complex formation. Schematic representation of the BRET setup is shown in K. L–M Correlation between the extent of endosomal β-arrestin2 translocation and MEK1 or ERK2 recruitment to AT_1_R, AUC values are shown, the same agonist concentrations were applied as in C, *N* = 4. Linear regression, r^2^ = 0.888, ***, P = 0.0001 for L, r^2^ = 0.8812, ***, P = 0.0002 for M. The corresponding kinetic curves are shown in Fig. 2C and Fig. S10A–B. Data are mean ± SEM in B, C, G L, and M.

Agonist-dependent endosomal β-arrestin2 recruitment was also visualized by confocal microscopy. First, the formation of intracellular β-arrestin2–Venus-enriched vesicles was assessed in live cells after stimulation with AngII, ST-AngII, or SII-AngII, which agonists have markedly different efficacies in β-arrestin recruitment (**Fig. 2F**). For the unbiased and high-throughput detection of intracellular fluorescent puncta, an unsupervised machine learning-based algorithm was applied **(Fig. S9)**. Significant differences were observed in the abilities of these ligands to form β-arrestin2–Venus-enriched vesicles (**Fig. 2G**). Administration of AngII led to the formation of a higher number of β-arrestin2–Venus-enriched puncta than ST-AngII or SII-AngII, moreover, the average size of the AngII-induced vesicles was also significantly greater (**Fig. 2H**–**I**). In addition to live-cell imaging, we performed quantitative analysis on fixed cells with an increased sample size for the full set of agonists. It should be noted that cell fixation per se caused artificial intracellular aggregates of β-arrestin2–Venus even in unstimulated cells, however the ligand-specific effects were clearly detectable. Confocal microscopy revealed a highly similar rank order of the agonists as the bystander BRET assay (**Fig. 2J**). These results verified that the observed temporal bias in β-arrestin recruitment is intimately associated with a spatial bias as the ligands markedly differed regarding their ability to induce endosomal β-arrestin binding.

Endosomal β-arrestin translocation was suggested to play an important role in β-arrestin-dependent regulation of the mitogen-activated protein kinase (MAPK) signaling cascade (*31*). Therefore, we also investigated whether the extent of endosomal β-arrestin translocation is in relation to the amount of complex formation between AT_1_R, β-arrestin2, and members of the MAPK pathway using previously described BRET assays (*32*) (**Fig. 2K**). There was a high correlation between the extent of endosomal β-arrestin translocation and the complex formation with MEK1 or ERK2, representing a downstream relevance of the magnitude and location of β- arrestin binding (**Fig. 2L**–**M, Fig. S10**).

### Ligand dissociation rate governs the lifetime of AT_1_R–β-arrestin2 complexes primarily in endosomes

All tested AT_1_ receptor agonists induced β-arrestin2 binding with various kinetics and efficacy, moreover, their signals were differently affected by the inhibition of endocytosis. Our next goal was to explore which intrinsic characteristics of the ligands can underlie these differences. To quantify the effects of internalization upon the β-arrestin2 signal of the different agonists, we investigated the difference between their E_max_ values with or without Dyn-K44A. We found that the β-arrestin2 signal of more efficacious ligands is systematically less sensitive to internalization (**Fig. 3A**), suggesting that they are able to maintain a stable receptor–β-arrestin complex even after receptor trafficking to intracellular compartments. To directly investigate the stability of the AT_1_R–β-arrestin2 interaction, we characterized the disassembly of the AT_1_R–β-arrestin2 complex. We followed the dissociation of β-arrestin2–Venus from AT_1_R–RLuc after the termination of agonist binding, which was achieved by ligand displacement with the high-affinity AT_1_R antagonist, candesartan (**Fig. S11**). The rate of receptor–β-arrestin disassembly (k_dis_) was assessed by using the exponential decay equation. Similar to the “internalization sensitivity”, the rate of β-arrestin2 detachment from AT_1_R greatly varied between different agonists and displayed a strong inverse correlation with their efficacy values (**Fig. 3B**).

**Figure 3.**
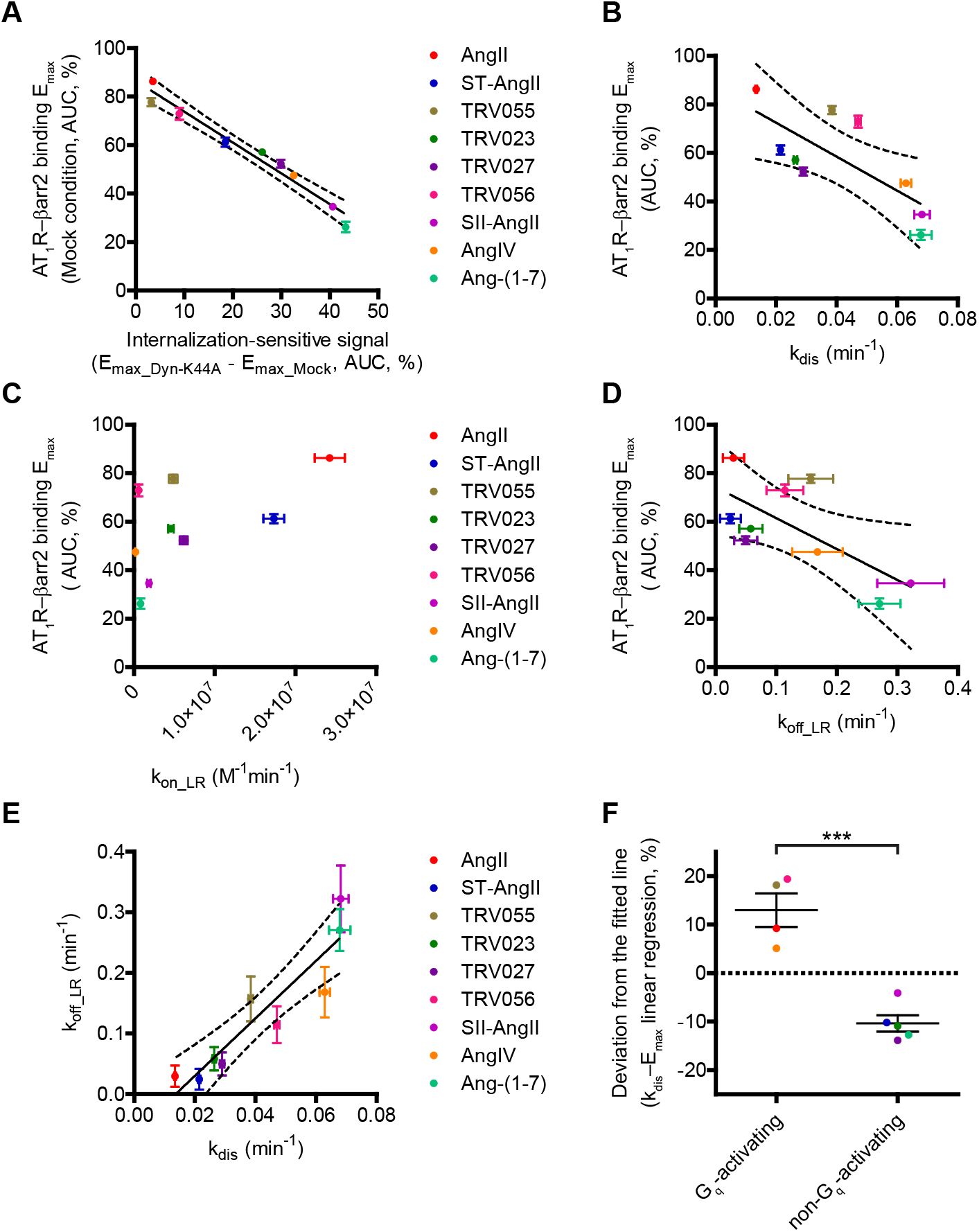
Ligand dissociation rate inversely correlates with the efficacy of AT_1_R–β-arrestin2 interaction. **A** Internalization sensitivity of AT_1_R–β-arrestin2 binding inversely correlates with its efficacy. The efficacy and degree of internalization sensitivity of each ligand were evaluated from concentration–response curves from Fig.1E. Internalization-sensitivity was quantified by the difference of E_max_ values in the presence or absence of Dyn-K44A. Linear regression, dotted lines represent 95% confidence intervals, r^2^ = 0.9643, ****, P < 0.0001. **B** Dissociation rate of β-arrestin2–Venus from AT_1_R–RLuc (k_dis_) inversely correlates with the efficacy of AT_1_R–β-arrestin2 binding (r^2^ = 0.5476, *, P = 0.0226). Kinetic curves are shown in Fig. S11. **C**–**D** Correlation between E_max_ of AT_1_R–β-arrestin2 binding and k_on_LR_ (C) or k_off_LR_ (D). K_off_LR_ showed a significant inverse correlation (r^2^ = 0.4794, *, P = 0.0387), while k_on_LR_ did not correlate with the extent of β-arrestin2 binding (r^2^ = 0.3637, n.s., P = 0.0982). **E** High degree of correlation between k_dis_ and k_off_LR_ (r^2^ = 0.8523, ***, P = 0.0004). **F** G_q_-activating ligands had significantly higher β-arrestin2 binding efficacy compared to non-G_q_- activating ligands, irrespectively from their corresponding k_dis_. Deviation from the fitted line in Fig. 3B is plotted for each ligand. Unpaired t-test was performed, ***, P = 0.0003. Data are mean or mean ± SEM.

Since the observed k_dis_ values incorporate the dissociation rates of complex reaction steps between agonists, receptors, and β-arrestin molecules, we tested whether the marked differences between ligands are driven by their kinetic binding parameters. We performed competitive ligand binding measurements to assess their receptor association and dissociation rates (k_on_LR_ and k_off_LR_ values), using a *Gaussia* luciferase (GLuc)-based BRET platform, we have described recently (*33*) (**Fig. S12**). All calculated values are shown in Table 1. While no significant correlation was found between k_on_LR_ and the extent of AT_1_R–β-arrestin2 binding, k_off_LR_ showed a similar significant inverse correlation as k_dis_ with the efficacy of the AT_1_R–β-arrestin2 interaction (**Fig. 3C** and **D**). Moreover, the k_off_LR_ values, obtained from the direct GLuc-based ligand binding assay, highly correlated with the k_dis_ values of the β-arrestin2 dissociation assay (**Fig. 3E**). These suggest that the dissociation rate of ligands is a major contributor to the agonist-dependent differences.

**Table 1.** Ligand association and dissociation rates determined by the GLuc-based competitive ligand binding assay

| | $k_{on\_LR} (M^{-1} min^{-1})$ | | $k_{off\_LR} (min^{-1})$ | |
| --- | --- | --- | --- | --- |
|  | Mean | S.E.M | Mean | S.E.M |
| AngII | $2.42 \times 10^7$ | $1.86 \times 10^6$ | $2.96 \times 10^{-2}$ | $1.74 \times 10^{-2}$ |
| ST-AngII | $1.73 \times 10^7$ | $1.28 \times 10^6$ | $2.46 \times 10^{-2}$ | $1.72 \times 10^{-2}$ |
| TRV055 | $4.86 \times 10^6$ | $5.41 \times 10^5$ | $1.57 \times 10^{-1}$ | $3.70 \times 10^{-2}$ |
| TRV023 | $4.57 \times 10^6$ | $3.44 \times 10^5$ | $5.83 \times 10^{-2}$ | $1.91 \times 10^{-2}$ |
| TRV027 | $6.15 \times 10^6$ | $4.80 \times 10^5$ | $4.99 \times 10^{-2}$ | $1.89 \times 10^{-2}$ |
| TRV056 | $5.75 \times 10^5$ | $5.87 \times 10^4$ | $1.15 \times 10^{-1}$ | $3.04 \times 10^{-2}$ |
| SII-AngII | $1.85 \times 10^6$ | $2.14 \times 10^5$ | $3.22 \times 10^{-1}$ | $5.51 \times 10^{-2}$ |
| AngIV | $1.77 \times 10^5$ | $2.17 \times 10^4$ | $1.68 \times 10^{-1}$ | $4.16 \times 10^{-2}$ |
| Ang-(1-7) | $8.14 \times 10^5$ | $6.54 \times 10^4$ | $2.71 \times 10^{-1}$ | $3.45 \times 10^{-2}$ |

### G protein activity regulates β-arrestin2 signaling in an endocytosis-dependent manner

Our previous analysis showed an overall strong correlation between ligand dissociation rate and β-arrestin2 binding efficacy, however G_q_-activating ligands generally displayed higher efficacy than biased agonists, independently from their k_off_LR_ and k_dis_ values (**Fig. 3F**). This is well illustrated by the examples of the balanced agonist AngII and the β-arrestin-biased ST-AngII (**Fig. 4A**). These ligands share almost the same k_off_LR_, but their efficacies for β-arrestin binding substantially differ, suggesting that G protein-dependent regulatory factors play an important role in the spatiotemporal control of β-arrestin binding. To address this question, we applied genetic and pharmacological perturbations. We first investigated the effects of a complete blockade of G protein activity using a G protein knockout CRISPR/Cas9 cell line (ΔGsix: ΔG_s/olf_/ΔG_q/11_/ΔG_12/13_), expressing only the G_i/o_ subfamily (*34*), which were pretreated with the G_i/o_-inhibitor pertussis toxin (PTX). In these cells, the differences between the AngII and ST-AngII effects completely disappeared, and the extent of β-arrestin2 binding to AT_1_R was also greatly reduced (**Fig. 4B**–**C**). We tested whether the G protein-mediated effects are under the spatiotemporal regulation of receptor trafficking. Remarkably, the inhibition of endocytosis with Dyn-K44A co-expression partially restored the lower β-arrestin binding in ΔGsix cells (**Fig. 4B**–**C**), suggesting that G proteins play an important role in the modulation of endosomal β-arrestin2 recruitment. In agreement with that, quantitative confocal microscopic experiments revealed significantly less β- arrestin2–Venus-enriched intracellular vesicles in AngII-stimulated ΔGsix cells (**Fig. 4D**–**E**).

**Figure 4.**
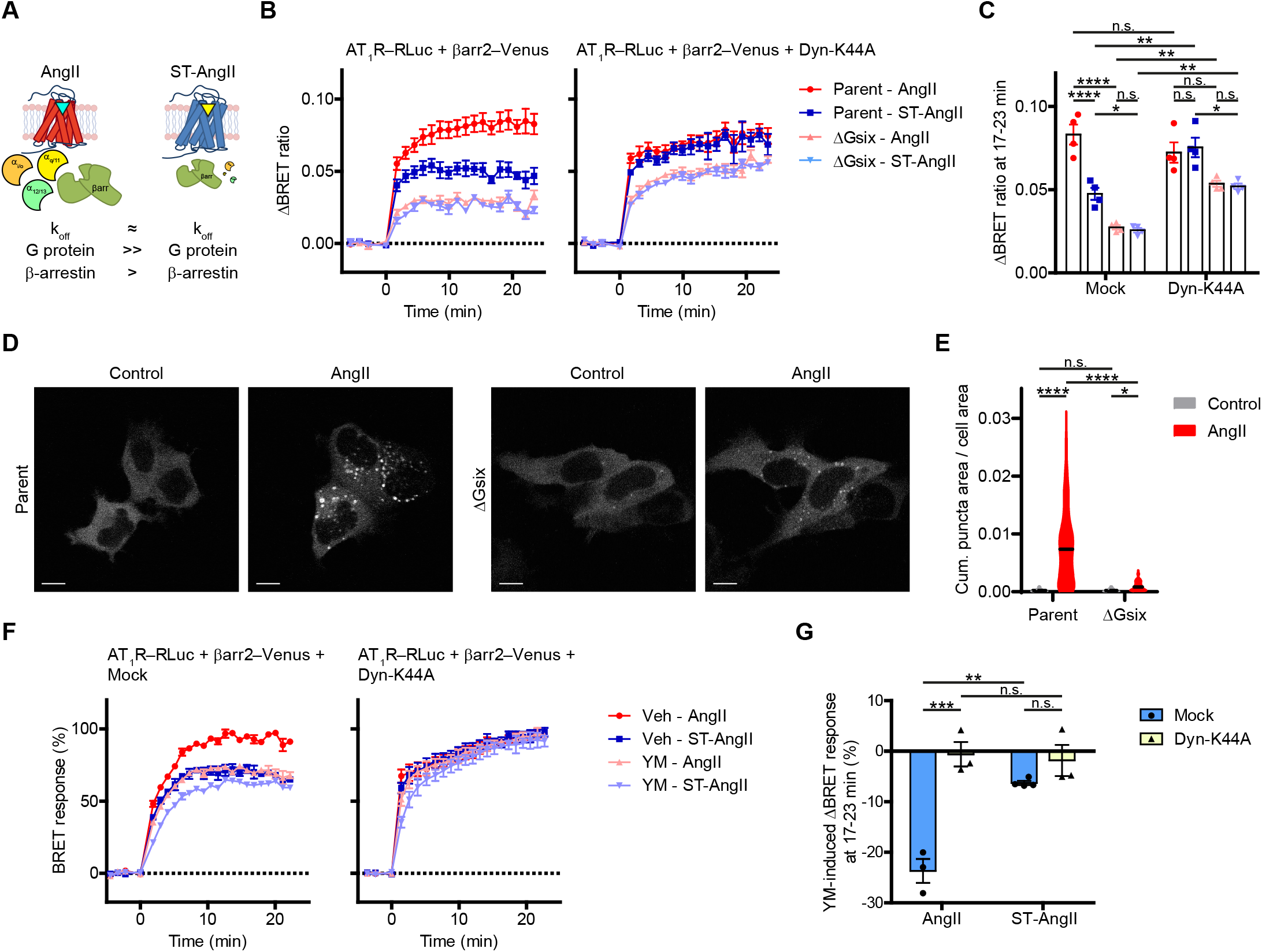
G protein activity maintains the endosomal β-arrestin2 recruitment to AT_1_R. **A** Schematic comparison of the characteristics of AngII and ST-AngII, representative high-affinity members of the G_q_- and non-G_q_-cluster agonists. **B** Kinetics of β-arrestin2–Venus binding to AT_1_R–RLuc in parent and PTX-pretreated (100 ng/ml, 20 h) ΔGsix HEK 293A cells after treatment with 10 μM AngII or ST-AngII. The absence of G protein activity completely abolished the agonist-dependent differences in β-arrestin2 signaling and reduced the effects of both drugs (left panel). However, the smaller extent of AT_1_R–β-arrestin2 binding in ΔGsix cells is rescued upon inhibition of receptor endocytosis by Dyn-K44A co-expression (right panel). **C** Statistical comparison of the changes in BRET ratios at later timepoints (17–23 min after stimulation) using three-way ANOVA with Bonferroni post-hoc test. Only the biologically meaningful comparisons are shown. The changes in the BRET ratios and the differences between ligand-induced effects were greatly decreased in ΔGsix cells. The reduced AT_1_R–β-arrestin2 binding in ΔGsix cells is significantly elevated if receptor endocytosis is blocked. **D** Representative live cell images of parent and PTX-pretreated ΔGsix HEK 293A cells before and after stimulation with 10 μM AngII. Scale bars are 10 μm, *N* = 4. **E** Quantitative analysis of live cell images of untreated and 20–30 min stimulated cells. 90, 194, 301, and 396 cells with similar β-arrestin2–Venus expression were analyzed for each conditions. Outliers were identified and excluded from the data using the ROUT method (Q = 1%). **F** Kinetics of β-arrestin2–Venus recruitment to AT_1_R–RLuc in vehicle or YM-254890 (YM, 100 nM, for 40 min)-pretreated HEK 293T cells. The G_q/11_-specific inhibitor YM decreased the maximal BRET responses, and it had a stronger effect on the G_q_-activator AngII in comparison to the non-G_q_-activator ST-AngII. YM effects were greatly rescued in Dyn-K44A co-expressing cells. **G** YM-related changes in the BRET response were abolished in internalization-inhibited condition at later time point (17–23 min after stimulation). YM-induced relative changes are shown, data are normalized to vehicle pretreatment for each ligand. Statistical evaluation was performed with two-way ANOVA with Bonferroni post-hoc test. Data are mean ± SEM, *N* = 3–4. N.s.: P >0.05; *, 0.05 ≥ P > 0.01; **, 0.01 ≥ P > 0.001; ***, 0.001 ≥ P > 0.0001, ****, P ≤ 0.0001.

To selectively evaluate the role of G_q/11_ protein activity, we conducted experiments with a specific G_q/11_-inhibitor, YM-254890 (YM) (*35*), after the verification that the drug effectively and selectively inhibits G_q/11_ proteins (**Fig. S13**). In the presence of YM, the AngII-induced response was markedly decreased, however its effect was weaker in the case of ST-AngII, in line with their different degree of G_q/11_ activation (**Fig. 4F**). Similarly as in ΔGsix cells, the inhibition of endocytosis diminished the effects of G_q/11_ inhibition on the agonist-induced β-arrestin2 binding curves (**Fig. 4F**–**G**). These implicate an important role of G protein activity in the maintenance of endosomal β-arrestin2 binding.

G_q/11_ activity also leads to PtdIns(4,5)P_2_ hydrolysis, which may decrease the extent of receptor internalization (*25*) and thus may further contribute to the enhanced β-arrestin2 binding. Notably, β-arrestin2 binding measurements require overexpression of β-arrestin2, which may influence plasma membrane PtdIns(4,5)P_2_ levels (*36, 37*). In the presence of overexpressed β- arrestin2, the agonist-induced PtdIns(4,5)P_2_ depletion was only transient **(Fig. S14A**–**B)**, and there was no substantial difference in the extent of receptor internalization induced by the distinct ligands **(Fig. S14C**–**F)**. These results also contradict that the differences in β-arrestin2 binding would be attributed to distinct ligand-induced internalization properties.

### A class B-type mutant β_2_-adrenergic receptor shows endocytosis-dependent, agonist-specific differences in β-arrestin recruitment

GPCRs are traditionally divided into two classes (A and B), based on their ability to maintain β- arrestin binding (*38*). AT_1_R belongs to class B receptors, which form a stable complex with β- arrestin2 at the endosomal compartment, where ligand-dependent differences in β-arrestin2 binding mostly emerged in our previous set of experiments. To address the question, if the observed findings are general among class B GPCRs, we extended our investigations to the β_2_- adrenergic receptor (β_2_AR), a prototypical class A GPCR, that is known to be incapable of endosomal β-arrestin recruitment. We hypothesized that the artificial induction of an endosomal pool of β_2_AR–β-arrestin2 complexes may augment differences between distinct β_2_AR agonists to recruit β-arrestin2.

To test this hypothesis, we utilized a mutant form of β_2_AR which is converted to a class B receptor by incorporation of C-terminal phosphorylation sites into its C terminus (β_2_AR-3S) (*39*), which can cause sustained β-arrestin2 binding at the endosomal compartment (**Fig. 5A**). In the case of wild type (WT) β_2_AR receptors, we found similar β-arrestin2 recruitments for the three tested β_2_AR agonists (**Fig. 5B**). However, the extent of interaction between the mutant β_2_AR-3S receptor and β-arrestin2 was not only elevated in general, but remarkable differences emerged between the effects of ligands (**Fig. 5C**), which did not occur in the case of the wild-type receptor. Moreover, if we inhibited receptor internalization with Dyn-K44A, these alterations were almost completely eliminated (**Fig. 5D**–**E** and **S15**), suggesting that ligand-dependent differences with the β_2_AR-3S receptor arose from the distinct ability of the β_2_AR agonists to induce endosomal β- arrestin binding.

**Figure 5.**
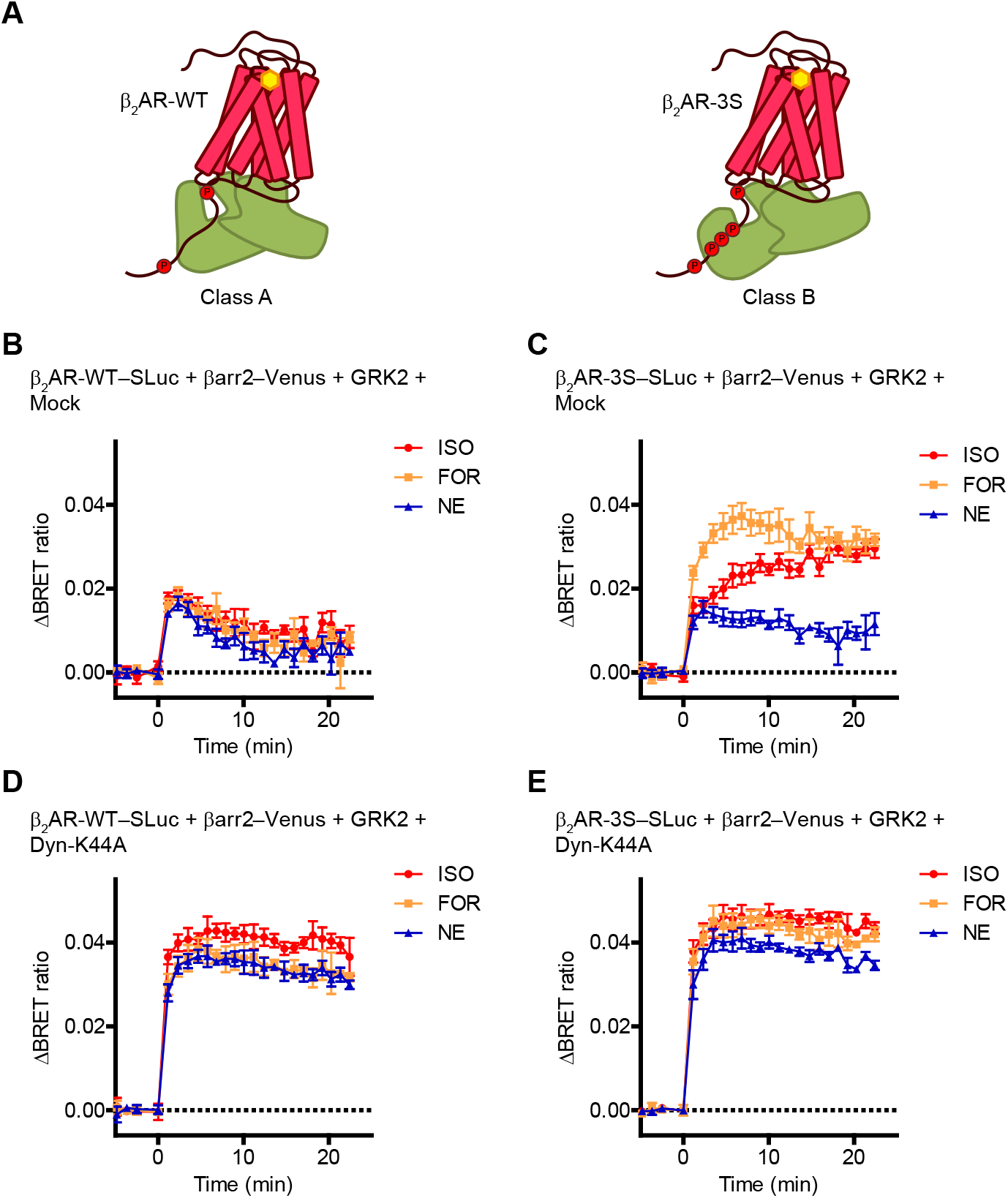
Artificial induction of endosomal β-arrestin binding of β_2_AR generates ligand-specific differences in β-arrestin2 recruitment. **A** Schematic representation of the β-arrestin binding properties of the wild-type (WT) and the phosphorylation site-engineered mutant (3S) β_2_AR. **B** Kinetics of the interaction between wild-type (WT) β_2_AR–SLuc, a prototypical class A receptor, and β-arrestin2–Venus. Formoterol (FOR, 30 μM), isoproterenol (ISO, 30 μM) and norepinephrine (NE, 300 μM) were used as stimuli, at concentrations that exert maximal β-arrestin2 binding. No differences in the BRET responses were found. **C** Stimulation of the β_2_AR-3S mutant receptor, which possess class B-type binding, resulted in higher β-arrestin2 signal and there was a substantial difference in the responses to FOR, ISO, and NE treatments, contrarily to the results obtained with wild-type β_2_AR. **D**–**E** Inhibition of endocytosis, using Dyn-K44A, significantly increased the extent of β-arrestin2 binding of both the wild-type and the 3S-mutant receptors. Moreover, in the case of β_2_AR-3S (E), it diminished the differences in agonist effects, suggesting that the various signals of the ligands were caused by their distinct ability to induce β-arrestin2 binding in endosomes. Corresponding concentration–response curves for B-E are shown in Fig. S15. The cells were transfected with the indicated constructs. Data are mean ± SEM, *N* = 3–4.

These results suggest that endocytosis may regulate the lifespan of agonist-induced GPCR- β-arrestin complexes and acts as a general orchestrator of the temporal effects of kinetic ligand parameters and dynamic system factors.

### Quantitative modeling reveals kinetic factors that regulate the endosomal β-arrestin recruitment

Our results with AT_1_R and β_2_AR-3S implicate that ligand-dependent differences in β-arrestin binding of class B GPCRs predominantly manifest in intracellular compartments. However, the precise experimental identification of the underlying internalization-sensitive molecular factors and their selective analyses faces technical limitations. To overcome these shortcomings and to investigate our concept with an independent approach, we constructed a kinetic mathematical model of GPCR signaling that allows the individual analysis of the relevant parameters in a compartment-specific manner.

We formulated ordinary differential equations (ODEs) to describe how the G protein activation and β-arrestin binding of receptors evolve over time upon agonist stimulation. Our complete modeling framework is displayed in **Fig. S16**, in which receptor internalization is also included. The reaction rate constants and the initial concentration of molecules were either chosen from previously introduced mathematical models of GPCR signaling and published experimental data or were set based on rational assumptions (**Table S1**–**S2**) (*40–44*). Our simulations were able to yield G_q_ activity and β-arrestin binding concentration response curves and to display the time-course of downstream signaling events mediated by G_q_ proteins (**Fig. S17**)

To investigate our experimental findings, we carried out simulations that examine the spatiotemporal aspects of β-arrestin binding and its relationship with the ligand dissociation rate constant (k_off_LR_). A well-known difference between the local regulation of GPCR signaling at endosomes is the relative rate of receptor phosphorylation and dephosphorylation compared to the plasma membrane (*45–48*). To model this, we set the receptor phosphorylation rate at the plasma membrane higher, and selectively evaluated the number of β-arrestin-bound receptors at the two different compartments. In agreement with our experimental results, a ligand with higher k_off_LR_ induced lower β-arrestin binding and displayed a different kinetic profile (**Fig. 6A**). In addition, the k_off_LR_-dependent differences were more prominent in the intracellular compartment (**Fig. 6B** and **C**). In agreement with these, the in silico inhibition of internalization greatly reduced the effects of the dissociation rate constant (**Fig. 6A**–**D**). These simulations reveal that the experimental correlation between k_off_LR_ and E_max_ implies a direct causation and confirm that the effects of k_off_LR_ are endocytosis dependent.

**Figure 6.**
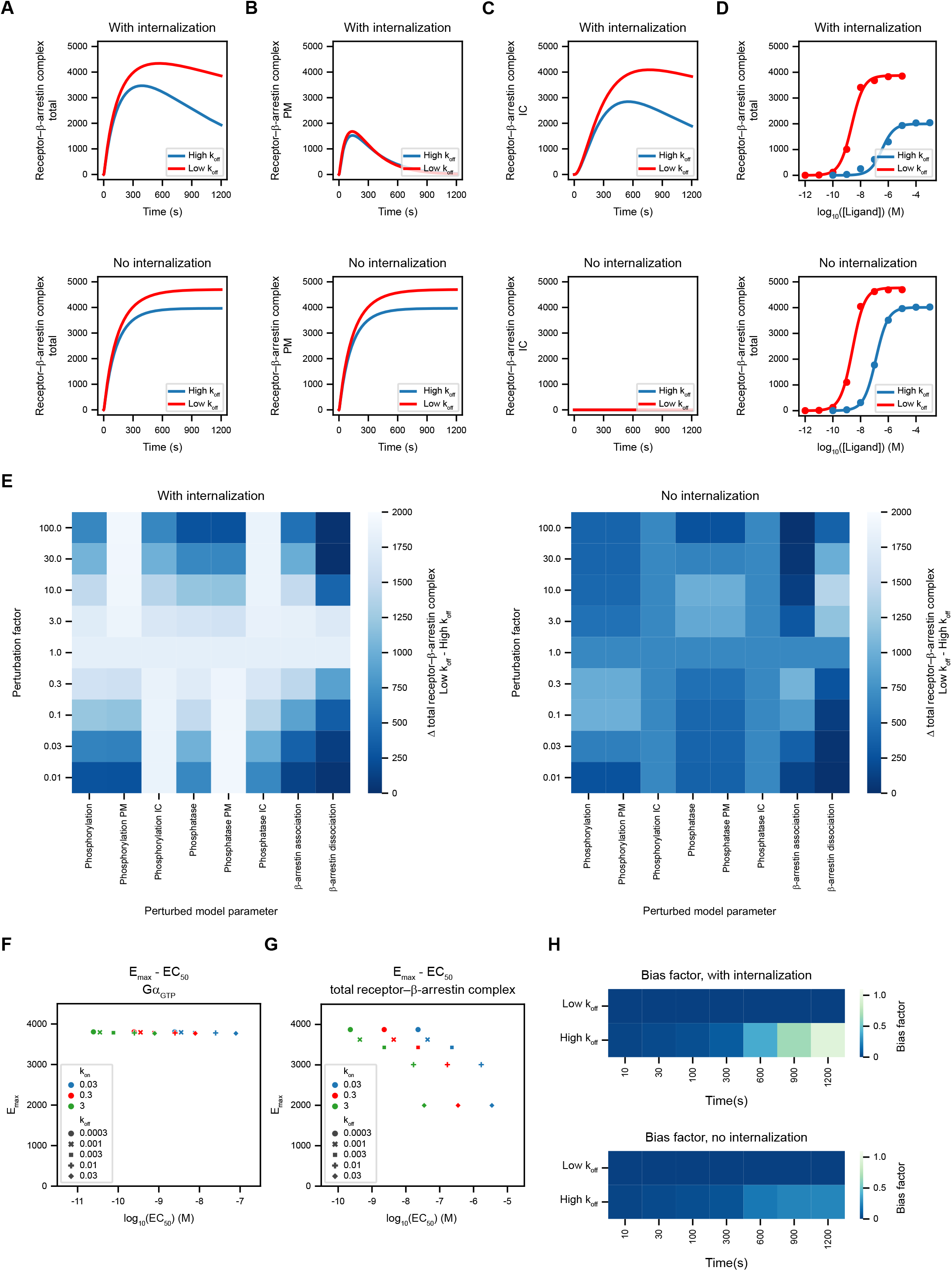
Quantitative kinetic modeling reveals mechanistic insights into spatiotemporal bias. **A–C** Simulated time course profiles of β-arrestin binding upon stimulation with two agonists, which only differ in their receptor dissociation rate constant (k_off_LR_: reactions 2, 4, 6, 8, 10, 12, 14, 16, 18, 20, 22, 24, 52, 54, 56, 58; “low k_off_”: k_off_LR_ = 0.0003, “high k_off_”: k_off_LR_ = 0.03). Top graphs represent simulated curves in the presence of receptor trafficking. Receptor internalization rate (reactions 38, 39, 40) was set to 0 to assess bottom graphs. A ligand with low k_off_LR_ resulted in the formation of more β-arrestin-bound receptors (**A**) and its higher effectiveness was more pronounced at the endosomal compartment (**C**) than at the plasma membrane (**B**) (total β-arrestin binding: molecules 10, 11, 12, 13, 14, 15; plasmalemmal β-arrestin binding: molecules 10, 11, 12; intracellular β-arrestin binding: molecules 13, 14, 15). The ligand-dependent differences between the two ligands were greatly reduced if endocytosis was blocked (bottom graphs). **D** Simulated concentration–response curves of the same agonists as in A–C. The total amount of β-arrestin-bound receptors at 20 min after stimulation are shown. Blockade of receptor internalization greatly reduced the difference between the efficacies of the ligands. **E** Effect of perturbation of reaction rate constants of the β-arrestin binding pathway. Simulations were performed by multiplying the initial rate constants of the investigated reactions with the indicated factors (Phosphorylation PM: reaction 30; Phosphorylation IC: reaction 50; Phosphorylation IC PM: reactions 30, 50; Phosphatase PM: reactions 25, 27, 29; Phosphatase IC: reactions 47, 48, 49; Phosphatase IC PM: reactions 25, 27, 29, 47, 48, 49; β-arrestin association: reactions 35, 36, 37, 44, 45, 46; β-arrestin dissociation: reactions 32, 33, 34, 41, 42, 43). The total amount of β-arrestin-bound receptors were assessed in systems with or without internalization for the same ligands as in A–D and the difference of 20-min values is plotted. The results for each ligand are shown in Fig. S18. K_off_-dependent differences substantially changed under each perturbed conditions, however the blockade of endocytosis decreased these effects. **F–G** Concentration–response curves were simulated for larger set of test ligands with different k_on_LR_ and k_off_LR_ values, which are indicated by the colors or the shape of symbols, respectively (k_on_LR_: reactions 1, 5, 9, 13, 17, 21, 51, 55). **G** Systematic increase of k_off_LR_ resulted in a graded decrease of the total amount receptor–β-arrestin, but perturbation of k_on_LR_ did not change the E_max_ values, only EC_50_ values were shifted. **F** The number of activated G proteins (molecules 23, 33) was not sensitive to any of the kinetic ligand binding parameters. **H** Heat maps visualize the time-dependence of the degree of G protein bias in the presence (top) or absence (bottom) of receptor trafficking. The extent of bias towards G protein activation vs. β- arrestin recruitment was quantified using the equiactive comparison equation *(24)*, and the low k_off_ agonist was set as reference ligand. The calculated bias factor displayed a marked temporal rise for the high k_off_ ligand. However, it remained “relatively balanced” even at later time points if endocytosis was inhibited.

β-arrestin binding is known to be modulated by numerous cellular regulatory mechanisms, that affect the phosphorylation state of receptors or change the affinity of the receptor–β-arrestin complex otherwise. To test whether such system factors can also influence the k_off_LR_-dependent effects, we perturbed the reaction rate constants of the β-arrestin binding pathway. The k_off_LR_- specific differences in β-arrestin recruitment were highly sensitive to changes in any of the investigated reaction rates (**Fig. 6E, Fig. S18**). Increased relative rate of phosphorylation at the endosomes, compared to the plasma membrane (either by decreasing plasma membrane phosphorylation / endosomal dephosphorylation or by increasing endosomal phosphorylation / plasma membrane dephosphorylation) diminished the k_off_LR_-specific differences. In the case of receptor–β-arrestin binding parameters, both increased and decreased stability of receptor–β- arrestin complex led to decreased k_off_LR_-specific differences. Moreover, we found that without receptor endocytosis the difference between a low and a high k_off_ ligand was generally less affected by our perturbations (**Fig. 6E, right**). These findings suggest that the modulation of the relationship between ligand k_off_LR_ and β-arrestin2 efficacy may serve as an important way to fine-tune signaling. On the other hand, changes in the ligand dissociation rate did not affect the maximal extent of G protein activation (**Fig. 6F**), suggesting that kinetic ligand parameters have disparate effects on distinct receptor-stimulated pathways. In contrast to k_off_LR_, alterations of ligand k_on_LR_ were not associated with marked changes in the efficacy of the investigated transducers (**Fig. 6F**– **G**).

To systematically analyze the role of k_off_LR_ in the apparent pathway selectivity, we ran simulations with a set of test agonists with gradually altered k_off_LR_ values and calculated a bias factor to quantify their relative preference towards G_q_ activation over β-arrestin binding (**Fig. 6H**). In the presence of receptor trafficking, k_off_LR_ emerged as a decisive attribute of ligands in their “functional selectivity”. However, without internalization, the calculated bias remained almost completely unaltered by k_off_LR_, further highlighting the role of compartmentalization in functionally selective signaling.

## DISCUSSION

In this study, we demonstrate that functionally selective signaling of GPCRs is a concerted interplay of intrinsic characteristics of the agonist-activated receptor structure (ligand bias), kinetic parameters (temporal bias), and spatial factors (location bias), that are strongly connected and strictly coordinated by the phenomenon of receptor endocytosis. We applied a diverse set of experimental and in silico approaches to unveil how receptor trafficking organizes these “types of bias” and display our results with AT_1_R, a prototypical GPCR with the capability of biased signaling. We found that inhibition of receptor internalization eliminates the differences between the AT_1_R–β-arrestin binding efficacies of distinct agonists, including classically considered biased and balanced ligands. We provide a mechanistic insight behind the profound effects of receptor trafficking by unveiling that ligand-dependent regulatory factors of β-arrestin binding are mainly exerted at the endosomal compartment.

Despite great research progress regarding “compartmentalized signaling” and “biased signaling”, the molecular links between these two phenomena remain poorly understood. On the other hand, their joint translational potential was recently highlighted by Eiger *et al*., who demonstrated that endocytosis is necessary for a β-arrestin-biased agonist to exert its anti-inflammatory effects in mice (*23*). These results implicate the unexploited possibility to rationally design biased drugs with defined spatiotemporal pharmacological profiles. Our experimental work addressed this concept and managed to identify the principal characteristics of ligands that determine signaling efficacy and the extent of bias in a compartment-specific manner.

First, we found that higher ligand dissociation rate is linked with faster disassembly of the agonist–receptor–β-arrestin complexes and thus decreases the total amount of β-arrestin-bound receptors. Our results regarding the strong influence of ligand dissociation rate constant on the overall β-arrestin binding efficacy are supported by previous observations made with other GPCRs (*19, 49, 50*). However, a recent study by Mösslein *et al.* contradicted this hypothesis since they found no effect of k_off_ on receptor–β-arrestin complex stability during the investigation of β_2_- adrenergic and μ-opioid receptors (*51*), which possess class A-type β-arrestin binding, i.e. are not able to recruit β-arrestin in endosomes. In this study, we resolve this apparent discrepancy, since we not only verified that k_off_LR_ regulates the amount of receptor–β-arrestin complexes, but also demonstrated that mainly the endosomal pool is affected by this ligand kinetic parameter. In agreement with the study of Mösslein et al., we found no differences between the efficacy of low-and high-affinity agonists with the wild-type β_2_AR, except for a phosphorylation site-engineered mutant that has class B-type β-arrestin binding properties. These results also draw attention to the fact that commonly applied signal-amplification solutions that transform the interaction type to class B, such as the C-terminal fusion of the C-tail of V_2_ vasopressin receptor (*52, 53*), may not only boost the extent of β-arrestin recruitment but artificially amplify differences between ligand efficacy values. This should be considered during the calculations of bias factors as well. In addition, our systematic in silico model suggests that the inhibition of internalization may increase the detected β-arrestin signal and thus may help to identify agonists with high k_off_ during ligand screening experiments.

Secondly, we found that endocytosis is a prerequisite for balanced agonists to induce a higher amount of receptor–β-arrestin interaction than β-arrestin-biased agonists via G protein activation. The inhibition of receptor trafficking did not alter ligand bias per se, and the G protein activity profiles of all the tested biased and balanced AT_1_R agonists were systematically resistant to the inhibition of endocytosis. In striking contrast, we found that the regulation of AT_1_R–β- arrestin interaction by the G protein activity of non-biased ligands heavily depends on receptor internalization. G protein activation greatly increased the degree of β-arrestin binding and this effect is even more pronounced in endosomes. The underlying molecular mechanism of this may be that G_q_- vs non-G_q_-activating agonists can engage different sets of GRK proteins (*7*). It is tempting to speculate that G proteins may directly activate GRKs in endosomes as well, however, at this time its selective experimental investigation faces unresolved technical difficulties.

Which special properties of the endosomal compartment can contribute to the locally different regulation of β-arrestin recruitment and how do they connect ligand characteristics with location bias? Our modeling approach highlighted that the relative activity of phosphorylation/dephosphorylation mechanisms is a possible main determinant of the magnitude of β-arrestin binding. Several observations of the current and previous studies implicate that dephosphorylation mechanisms dominate in endosomes due to relatively higher phosphatase and lower GRK activity (*46–48*), in contrast to the plasma membrane, where the high abundance of different GRK isoforms strongly shifts the regulatory reactions in favor of receptor phosphorylation (*54*). Furthermore, not only the quantity of phosphorylated receptors matters, but the sequence of the phosphorylation residues, also known as the phosphorylation barcode, has equally important effects on the extent, kinetics, and conformation of β-arrestin binding. Recent evidence suggests that distinct ligands have different phosphorylation barcodes, which manifest in functionally selective signaling outcomes (*55–57*). Moreover, ligands may be biased regarding their endosomal GRK activation profiles, which further contributes to “location bias” (*58*).

The distinct lipid composition of intracellular membranes, such as the absence of PtdIns(4,5)P_2_, adds another layer of complexity to the endosomal regulation of signal transduction. The high level of plasmalemmal PtdIns(4,5)P_2_ was recently suggested to play a key role in stabilizing the active state of agonist-bound receptors and modulating the receptor–β-arrestin complex formation (*59, 60*). Conversely, the lack of PtdIns(4,5)P_2_ in endosomal membranes may facilitate the disassembly of agonist–receptor–β-arrestin complexes.

Compartment-specific characteristics of the ligand–receptor interaction can also distinguish endosomal β-arrestin binding from that in the plasma membrane. The relatively acidic environment of endosomes may accelerate ligand dissociation (*61*) and the consequent lower receptor residence time can promote receptor resensitization (*62*). Moreover, the luminal ligand concentrations in different subcellular compartments may also differ from that in the extracellular space. Due to the relatively small volume of endosomes, ligand concentrations can be even higher. However, ligand depletion may also occur in endosomes, where endothelin converting enzyme 1 (ECE1) and other peptidases can rapidly cleave peptide ligands, which prevents rebinding and thus shortens the signal transmission of intracellular receptors (*63, 64*). Since the kinetic constants that describe changes in endosomal pH or ligand concentrations are not precisely known, these were not incorporated into our mathematical model. Nevertheless, one can speculate that including these mechanisms could further strengthen the conclusion of our study.

We believe that our results can greatly assist the development of novel biased pharmaceutical compounds by improving our understanding of the molecular link between ligand characteristics and functional selectivity. A direct implication of this study is that higher endosomal β-arrestin recruitment is expected from ligands with long residence time, and total β- arrestin recruitment can be augmented by strategies that interfere with receptor internalization. Furthermore, our data propose that structure–activity relationship studies may highly benefit from the conduction of cell-based signaling assays both with and without the inhibition of receptor endocytosis. Experimental results under these two conditions could provide complementary information about the mechanism of novel drugs and help to separate the extent of ligand bias from other spatiotemporal factors which influence the pathway-specific efficacy of drugs.

## MATERIALS AND METHODS

### Compounds

TRV120023, TRV120027, TRV120055, TRV120056 (*24*), and TAMRA-AngII were synthesized by Proteogenix. [Sar^1^, Ile^1^, Ile^8^]-angiotensin II (SII-AngII) was from Bachem. YM-254890 was purchased from Wako Chemicals. Candesartan and formoterol were from Tocris. Rapamycin was bought from Selleckchem. Prolume Purple was obtained from NanoLight. Coelenterazine h was purchased from Regis Technologies. All other reagents were from Sigma-Aldrich.

### Plasmid constructs

The following constructs have been previously described: AT_1_R, AT_1_R–Rluc8, β_2_AR-3S–SLuc, β-arrestin2-K2A–Venus, Venus–Rab5 (*32*), AT_1_R–Rluc (*25*), β-arrestin1–Venus, β-arrestin2– Venus (*65*), L10–Venus (Venus fused to ‘L10’, the 10 first amino acids of mouse Lck protein, functioning as a myristoylated-palmitoylated plasma membrane target sequence), L10–Cerulean, plasma membrane PtdIns(4,5)P_2_ level BRET biosensor (L10–Venus–T2A–PLCδ1PH–SLuc) (*66*), GLuc–PM, NanoLuc–PM (*33*), β_2_AR–SLuc (*29*). PM–NanoLuc and EE–NanoLuc were generated by PCR amplification of the coding sequence of NanoLuc with or without stop codon using NanoLuc–PM as a template and replacing the Venus sequence with them in L10–Venus and Venus–Rab5, respectively (in PM–NanoLuc, the L10 sequence represents the target signal, whereas Rab5 protein marks early endosomes for EE-NanoLuc). PtdIns(4,5)P_2_ depletion system L10–FRB–T2A–FKBP–5-ptase construct was created by replacing the PM2 sequence with the L10 sequence with PCR amplification using the PM2–FRB–T2A–mRFP–FKBP–5-ptase as a template, then FKBP–5-ptase was fused in frame to the T2A sequence by replacing mRFP–FKBP– 5-ptase. TRUPATH was a gift from Bryan Roth (Addgene kit #1000000163) (*27*). Untagged β- arrestin2, GRK2, and HA–dynamin2A-K44A were kindly provided by Dr. Stephen S. Ferguson and Dr. Kazuhisa Nakayama. Venus–MEK1–FLAG and FLAG–ERK2–Venus constructs were kind gifts from Dr. Attila Reményi.

### Cell culture and transfection

The generation of HEK 293A ΔGsix (ΔG_s_/ΔG_olf_/ΔG_q_/ΔG_11_/ΔG_12_/ΔG_13_) cells was described previously (*34*). HEK 293T, HEK 293A parent and ΔGsix cells were cultured in DMEM supplemented with 10% FBS and 1% penicillin/streptomycin. The cells were transfected with the calcium phosphate precipitation method (in suspension for GLuc BRET measurements or on adherent cells for confocal microscopy measurements), or with Lipofectamine 2000 (in suspension, used for all other measurements) as previously described (*32, 33*). The plasmid DNA amounts applied are shown in Supplementary Table 3. For BRET measurements, transfected cells were cultured on white 96-well poly-L-lysine-coated plates. The BRET measurements with HEK 293A parent and ΔGsix cells were performed 48 h after transfection, in all other cases the experiments were made 24-28 h after transfection.

### Bioluminescence resonance energy transfer measurements

BRET measurements were executed using Thermo Fisher Varioskan or Varioskan Lux multimode plate readers similarly as previously described (*32, 33*). 24-28 hours after transfection, the media were replaced with a modified Krebs–Ringer solution (120 mm NaCl, 10 mm Na-HEPES, 10 mm glucose, 4.7 mm KCl, 1.2 mm CaCl_2_, 0.7 mm MgSO_4_, pH 7.4) including a washing step. The expression of fluorescent protein-tagged constructs was checked by fluorescence intensity measurements (emission at 535 nm wavelength with excitation at 510 nm for Venus, emission at 515 nm with excitation at 400 nm for GFP2 fluorescence). The used luciferase substrates filters are summarized in Supplementary Table 4.

The BRET measurements were performed at 37 °C, except the ligand binding measurements, which were made at 27 °C.

In kinetic measurements, first the basal BRET ratios were determined after the addition of the BRET substrate, then the indicated ligands were added, and BRET was followed continuously.

Basal BRET ratios were subtracted, and agonist-induced BRET changes were calculated by subtracting the BRET ratio of vehicle-treated cells. Unless otherwise stated, data were presented as a percentage of the AngII (100 nM or 10 μM)-induced change in the BRET ratio (BRET response). As coexpression of an additional protein to the BRET pair may influence the BRET ratio by altering the BRET donor/acceptor ratio, percentage expression was only used for the same expression conditions.

Plasma membrane PtdIns(4,5)P_2_ depletion was achieved by 300 nM rapamycin treatment for 5 minutes in cells expressing the PtdIns(4,5)P_2_ depletion system. Rapamycin induces the heterodimerization between the FRB and the FKBP domains of these constructs and thus leads to plasma membrane recruitment of the 5-phosphatase, which degrades PtdIns(4,5)P_2_ (*29*).

Competitive ligand binding measurements were performed in live cells similarly as described previously (*33*). Cells were co-expressed with AT_1_R, GLuc–PM, HA–dynamin2A-K44A, and β- arrestin2 constructs. To investigate the receptor occupancy of TAMRA–AngII, cells were treated with increasing concentrations of TAMRA–AngII for 2 hours at room temperature. Non-specific binding was assessed in cells cotreated with 10 μM candesartan, a high-affinity AT_1_R antagonist. Specific binding was determined by subtracting the non-specific signal from the total signal. Two-site specific binding curve was fitted to obtain the K_D_low_ and K_D_high_ values.

Thereafter, the kinetic rate constants (k_off_LR_ and k_on_LR_ values) of the interaction between TAMRA–AngII and AT_1_R were determined. To assess k_off_LR_, 1 μM TAMRA–AngII treatment was applied for 15 minutes, thereafter TAMRA–AngII was washed out, and media containing the BRET substrate and 10 μM candesartan (for prevention of rebinding) was added. The BRET ratio in time point 0 was determined in cells which were re-treated with TAMRA–AngII without candesartan. The data were normalized to the BRET ratio of cells which were not treated with AT_1_R ligands. The basal BRET ratios were not determined in these experiments. Two phase decay curve was fitted, and the initial proportion of the high-and low-affinity binding was calculated and set based on the previously fitted K_D_high_ and B_max_high_ values. As the high-affinity binding site is occupied mostly in the applied concentration, the k_off_LR_ value of the high-affinity binding site was used in further calculations. In k_on_LR_ measurements, after assessment of the basal BRET ratios, cells were treated with 300 nM TAMRA–AngII with or without 10 μM candesartan. The candesartan co-treatment was applied to determine the non-specific signal, which was subtracted from the total signal. One-site association binding curve (one conc. of hot) equation was used to calculate the k_on_LR_ value of TAMRA–AngII. The kinetic binding parameters of unlabeled AT_1_R ligands were assessed by following the BRET ratio change after simultaneous treatment of 1 μM TAMRA–AngII and increasing concentrations of the unlabeled ligands. For the calculation of k_on_LR_ and k_off_LR_ values of unlabeled ligands, we applied certain simplifications due to the large number of variables. We used a one-site binding model since in the applied TAMRA–AngII concentration mostly the high-affinity binding site is occupied as it is shown in Fig. S12B. In addition, we made the presumption that all agonist-bound receptors induce β-arrestin2 binding (ternary complexes) and ignored the ligand-occupied receptor state that is not coupled to β-arrestin. Kinetic binding parameters were fitted using the Motulsky–Mahan (kinetics of competitive binding) equation (*67*).

To assess the dissociation rate of β-arrestin2–Venus from AT1R–RLuc, first the AT_1_R–β-arrestin2 binding was induced by 12-min agonist treatment. The agonists were applied in ∼30 × EC_50_ concentrations. Thereafter, agonists were displaced by the addition of 10 μM candesartan, a high-affinity AT_1_R antagonist. One-phase dissociation curves were fitted to calculate the observed dissociation rate of β-arrestin2–Venus from AT_1_R–RLuc (k_dis_).

### Confocal fluorescence microscopy

For confocal microscopy imaging of fixed cells, cells were seeded on IBIDI µ-Slide 8 well plates coated with poly-L-lysine on the day before transfection. For the imaging of live cells, cells were seeded on poly-L-lysine-coated glass cover plates.

The adherent cells were cotransfected with plasma membrane-targeted Cerulean (L10–Cerulean), unlabeled AT_1_R, and β-arrestin2–Venus. In fixed cell experiments, cells were treated with AT_1_R ligands for 30 minutes in a modified Krebs–Ringer solution at 37 °C. Next, the cells were fixed with ice-cold 4% paraformaldehyde in phosphate buffered solution (PBS) for 15 minutes. Thereafter, the cells were washed three times with PBS for 5 minutes at room temperature.

For live cell experiments, coverslips were placed into a chamber, media was replaced with modified Krebs–Ringer solution and the measurements were performed at 37 °C. Fluorescence imaging was performed with a Zeiss LSM 710 confocal laser-scanning microscope using a 40× objective in tile scan mode (4×4) using autofocusing, with 2 µm offset from the well bottom.

### Image analysis

Image analysis was performed using Python. The cells were detected using the Cellpose library (*68*) on the L10–Cerulean channel. The vesicles were detected using the TensorFlow implementation of the Pix2Pix model (*69, 70*) trained on manually labeled images from the experiments. The further analysis of the cell-and vesicle masks aligned with the original images were done using the Pandas library (*71*). β-arrestin2–Venus expression levels were slightly different in the parent and the ΔGsix HEK 293A cells. Therefore, in the experiments in which both cell lines were used, only the cells in the fluorescent range present in both samples (250-600) were used. The Python code used in the analysis can be found on GitHub at https://github.com/turugabor/cellAnalysis.git.

### Mathematical modeling

We developed a mathematical model of G protein-coupled receptor (GPCR) signaling that captures the impact of various factors on receptor–β-arrestin binding. The model is based on ordinary differential equations (ODEs) and comprises 43 molecular species and 96 reactions, such as enzymatic reactions, binding events, and compartment changes. The molecular concentrations and reaction rate constants of the model (Table S1 and Table S2, respectively) were obtained from literature sources. The time derivatives of molecular concentrations were calculated using the reaction equations (Table S2) and were integrated numerically. Initial parameters of the ODE model — in the absence of ligand stimulation — represent a steady state.

Our model is composed of four basic modules: receptor–β-arrestin interactions, heterotrimeric G protein interactions, PLC activation, and second messenger generation (Fig. S17). Additionally, the model considers three compartments: plasma membrane, cytosol, and intracellular vesicles.

The receptor–β-arrestin module incorporates ligand binding to receptors, receptor activation and deactivation, receptor phosphorylation and dephosphorylation, β-arrestin binding, and receptor internalization (Fig. S17A). Importantly, internalized receptors can maintain ligand and β-arrestin binding. We modeled internalization as a unidirectional reaction and did not include receptor recycling or degradation in the model. Notably, the phosphorylation rate of intracellular receptors is 100 times lower than the plasma-membrane receptors. We also incorporated the rational assumption that the agonist-activated, phosphorylated receptor exhibits a higher affinity for β- arrestin compared to its phosphorylated, inactive counterpart.

As our simulation is tailored to the G_q/11_ protein-coupled AT_1_ angiotensin receptor, the G protein and PLC modules of our model reflect the signaling of G_q/11_ heterotrimeric G protein, including receptor–G protein interaction, G protein dissociation and reassociation (Fig. S17B), and interaction between the α subunit and PLC (Fig. S17C). Additionally, our second messenger module incorporates PtdIns(4,5)P_2_ synthesis and PLC-induced cleavage of PtdIns(4,5)P_2_ into diacylglycerol (DAG) and inositol 1,4,5-trisphosphate (IP_3_) (Fig. S17D).

We used three different sets of initial values in the simulations, one resembling a receptor / β- arrestin overexpressing system ([Receptor] = 5000 molecule / μm^2^ [Arrestin] = 15000 molecule / μm^2^, [G protein] = 40 molecule / μm^2^), another resembling a receptor / G protein overexpression system ([Receptor] = 5000 molecule / μm^2^, [Arrestin] = 1000 molecule / μm^2^, [G protein] = 4000 molecule / μm^2^) and another resembling receptor overexpression system ([Receptor] = 5000 molecule / μm^2^, [Arrestin] = 1000 molecule / μm^2^, [G protein] = 40 molecule / μm^2^). These systems correspond to typical experimental systems for measuring β-arrestin and G protein activation or second messenger generation, respectively.

We used Python 3.8 to run the simulations and for data analysis. Scipy library (*72*) was used for numerical integration. The code to reproduce our results is available on GitHub at https://github.com/bence-szalai/gpcr-signaling-simulation.

### Statistical analysis

The figures of the experimental data were generated by GraphPad Prism 9 software. The variance of ligand effects was calculated as the squared deviation of the agonist-induced responses. Log(agonist) vs. response curves were fitted on the concentration–response data. The bottom was constrained to 0, Hill slope was set to 1 for the β-arrestin2 binding data. Correlations were analyzed by linear regression and Pearson test. Unpaired or paired two-tailed t tests were used to compare the means of two distributions. Variances of distributions were analyzed with F tests, analysis was performed on average normalized datasets in order to compare two groups with identical means. Multiple groups, based on the experimental setting, were analyzed using one-way ANOVA, two-way ANOVA, or three-way ANOVA. Bonferroni post-hoc test was used if multiple comparisons were performed. Unless otherwise stated, kinetic data were normalized to baseline (data points before stimulation). Time scales were adjusted to better indicate the time length between stimulation and the first stimulated measurement point. Data of time point 0 represents the data of the last time point before stimulation. The time of one cycle length was subtracted from the time between the last baseline and first stimulated points and was added to the time between the last two baseline points. When distributions of agonist-induced β-arrestin2–Venus puncta in each identified cell were analyzed, outliers were identified and excluded using the ROUT method (Q = 1%). The k_off_LR_ and k_on_LR_ values for TAMRA–AngII were calculated with the association kinetics (one ligand concentration) and dissociation kinetics equations, respectively. The k_off_LR_ and k_on_LR_ values for unlabeled agonists were calculated by the kinetics of competitive binding equation. The bias factor was calculated using the equiactive comparison model (equation (6) in (*24*)). Data are mean ± SEM. All experiments were independently performed at least three times. BRET measurements were made in duplicate or triplicate, with the exception of MEK1 and ERK2 complex formation BRET assays, where sextuplicates were used.

### Supplementary Materials

Figure S1. Schematic illustrations of the BRET assays for transducer activation and the peptide sequence of the AT_1_R agonists

Figure S2. Agonist-induced β-arrestin1 and β-arrestin2 binding profiles of AT_1_R are highly identical

Figure S3. Systematic assessment of the agonist-specific G protein activation profile of AT_1_R using the TRUPATH BRET sensor set

Figure S4. Kinetics of β-arrestin2–Venus binding to AT_1_R–RLuc upon stimulation with increasing concentrations of agonists

Figure S5. Inhibition of endocytosis markedly changes the efficacy but not the potency of AT_1_R agonists in β-arrestin2 binding measurements

Figure S6. Inhibition of receptor trafficking by PtdIns(4,5)P_2_ depletion recapitulates the effects of endocytosis inhibition on AT_1_R–β-arrestin2 interaction

Figure S7. Effect of endocytosis inhibition with hypertonic sucrose on the transducer activation profile of AT_1_R agonists

Figure S8. G_q/11_–PLCβ signaling pathway is not significantly affected by the blockade of endocytic processes

Figure S9. Unsupervised, machine learning-based method for the detection of endosomal translocation of β-arrestin2–Venus

Figure S10. Real-time monitoring of AT_1_R–β-arrestin2–MEK1/ERK2 complex formation with BRET

Figure S11. Monitoring the β-arrestin2 dissociation from AT_1_R after agonist displacement

Figure S12. BRET-based measurement of the kinetic rate constants of ligand–receptor interaction in cells overexpressing β-arrestin2

Figure S13. G_q/11_-inhibitor YM-254890 effectively and selectively reduced AngII-induced G_q_ protein activity

Figure S14. Lack of PtdIns(4,5)P_2_ cleavage-mediated inhibition of endocytosis in β-arrestin2 overexpressing cells

Figure S15. Agonist-and endocytosis-dependent differences in β-arrestin2 recruitment of a β_2_AR mutant with engineered phosphorylation sites

Figure S16. Schematic representation of the molecules and reactions incorporated into the ODE model

Figure S17. Simulated time course profiles of major GPCR signaling events

Figure S18. Effects of perturbation of reaction rate constants of the β-arrestin binding pathway

Table S1. Initial concentration values of modeled molecular species

Table S2. Reaction equations and rate constants of modeled reactions Table S3. Applied plasmid DNA amounts in the distinct BRET setups

Table S4. Table of applied luciferase substrates and filters in BRET measurements

## Acknowledgments

The technical assistance of Eszter Halász and Kata Szabolcsi is greatly appreciated.

## Author contributions

Conceptualization: ADT, BS, GT, LH

Methodology: ADT, BS, AB, AI, PV, GT

Investigation: ADT, BS, OTK, DG, SP, GT, LH

Visualization: ADT, BS, DG, SP

Funding acquisition: GT, LH

Supervision: GT, LH

Writing: ADT, BS, OTK, DG, SP, AB, PV, GT, LH

## Data and materials availability

All codes applied are openly available as stated in the Methods section. All datasets are available from the corresponding authors on reasonable request.

