## Supplementary Materials for "Receptor endocytosis orchestrates the spatiotemporal bias of β-arrestin signaling"

Supplementary Figures S1–S18
Supplementary Tables S1–S4

### Table of contents

Figure S1. Schematic illustrations of the BRET assays for transducer activation and the peptide sequence of the $AT_1R$ agonists ... 3

Figure S2. Agonist-induced β-arrestin1 and β-arrestin2 binding profiles of $AT_1R$ are highly identical ... 4

Figure S3. Systematic assessment of the agonist-specific G protein activation profile of $AT_1R$ using the TRUPATH BRET sensor set ... 5

Figure S4. Kinetics of β-arrestin2–Venus binding to $AT_1R$–RLuc upon stimulation with increasing concentrations of agonists ... 7

Figure S5. Inhibition of endocytosis markedly changes the efficacy but not the potency of $AT_1R$ agonists in β-arrestin2 binding measurements ... 9

Figure S6. Inhibition of receptor trafficking by PtdIns(4,5)$P_2$ depletion recapitulates the effects of endocytosis inhibition on $AT_1R$–β-arrestin2 interaction ... 10

Figure S7. Effect of endocytosis inhibition with hypertonic sucrose on the transducer activation profile of $AT_1R$ agonists ... 12

Figure S8. $G_{q/11}$–PLCβ signaling pathway is not significantly affected by the blockade of endocytic processes ... 14

Figure S9. Unsupervised, machine learning-based method for the detection of endosomal translocation of β-arrestin2–Venus ... 16

Figure S10. Real-time monitoring of $AT_1R$–β-arrestin2–MEK1/ERK2 complex formation with BRET ... 17

Figure S11. Monitoring the β-arrestin2 dissociation from $AT_1R$ after agonist displacement ... 18

Figure S12. BRET-based measurement of the kinetic rate constants of ligand–receptor interaction in cells overexpressing β-arrestin2 ... 19

Figure S13. $G_{q/11}$-inhibitor YM-254890 effectively and selectively reduced AngII-induced $G_q$ protein activity ... 21

Figure S14. Lack of PtdIns(4,5)$P_2$ cleavage-mediated inhibition of endocytosis in β-arrestin2 overexpressing cells 22

Figure S15. Agonist- and endocytosis-dependent differences in β-arrestin2 recruitment of a $\beta_2$AR mutant with engineered phosphorylation sites ... 24

Figure S16. Schematic representation of the molecules and reactions incorporated into the ODE model ... 25

Figure S17. Simulated time course profiles of major GPCR signaling events ... 27

Figure S18. Effects of perturbation of reaction rate constants of the β-arrestin binding pathway ... 28

Table S1. Initial concentration values of modeled molecular species. ... 29

Table S2. Reaction equations and rate constants of modeled reactions ... 30

Table S3. Applied plasmid DNA amounts in the distinct BRET setups ... 32

Table S4. Table of applied luciferase substrates and filters in BRET measurements ... 33

**A**

| | | 1 | 2 | 3 | 4 | 5 | 6 | 7 | 8 |
|---|---|---|---|---|---|---|---|---|---|
| AngII | | Asp | Arg | Val | Tyr | Ile | His | Pro | Phe |
| TRV055 | | | **Gly** | Val | Tyr | Ile | His | Pro | Phe |
| TRV056 | Asp | Arg | **Gly** | Val | Tyr | Ile | His | Pro | Phe |
| AngIV | | | | Val | Tyr | Ile | His | Pro | Phe |
| ST-AngII | | **Sar** | Arg | Val | Tyr | Ile | His | Pro | **Thr** |
| TRV023 | | **Sar** | Arg | Val | Tyr | **Lys** | His | Pro | **Ala-OH** |
| TRV027 | | **Sar** | Arg | Val | Tyr | Ile | His | Pro | **D-Ala-OH** |
| SII-AngII | | **Sar** | Arg | Val | **Ile** | Ile | His | Pro | **Ile** |
| Ang-(1-7) | | Asp | Arg | Val | Tyr | Ile | His | Pro | |

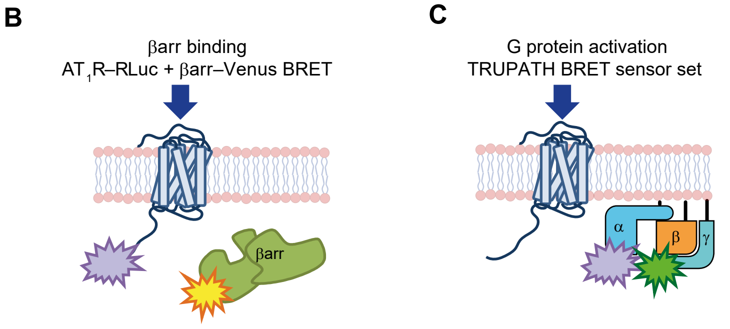


**Figure S1. Schematic illustrations of the BRET assays for transducer activation and the peptide sequence of the $AT_1R$ agonists**

**A** Amino acid sequence of the tested ligands in this study. β-arrestin-biased ligands (non-$G_q$-activating) are colored with blue, strong $G_q$-activator ligands are colored with red, the sequences are aligned to that of AngII.

**B** and **C**. Schematic representations of the BRET setups used for real-time monitoring of the transducer activation. Increase of the BRET ratio in the β-arrestin binding assays reflects the interaction between the RLuc-tagged receptor and Venus-tagged β-arrestin. In the TRUPATH system, the decrease of the BRET ratio reflects heterotrimeric G protein activation, when Rluc8-tagged Gα and GFP10-tagged Gγ subunits dissociate. In the figures, BRET response is displayed, which represents the change of BRET ratio (stimulated - unstimulated) as a percentage of the peak AngII (100 nM or 10 μM)-induced change of BRET ratio.

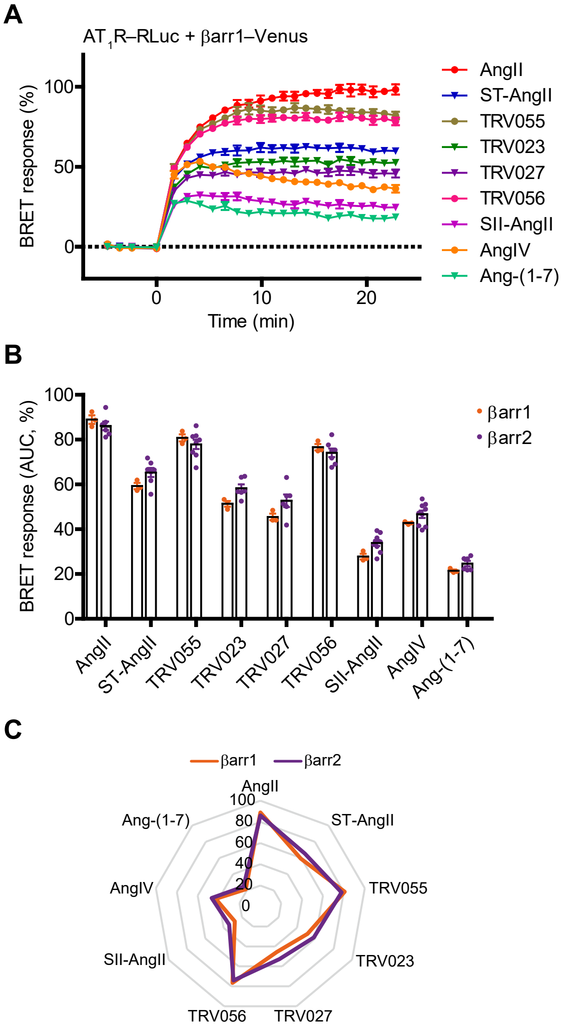


**Figure S2. Agonist-induced β-arrestin1 and β-arrestin2 binding profiles of $AT_1R$ are highly identical**

**A** Kinetics of the interaction between $AT_1R$–Rluc and β-arrestin1–Venus.

**B**–**C** Comparison of β-arrestin1–Venus and β-arrestin2–Venus binding to $AT_1R$–RLuc. All $AT_1R$ agonists were applied at 10 µM. Panel B shows scatter dot plot of AUC values, scatters represent independent biological replicates. Radial plot of mean values is displayed in C. Except panel C, data are mean ± SEM. $N = 3$ for β-arrestin1 and $N = 6$–$9$ for β-arrestin2.

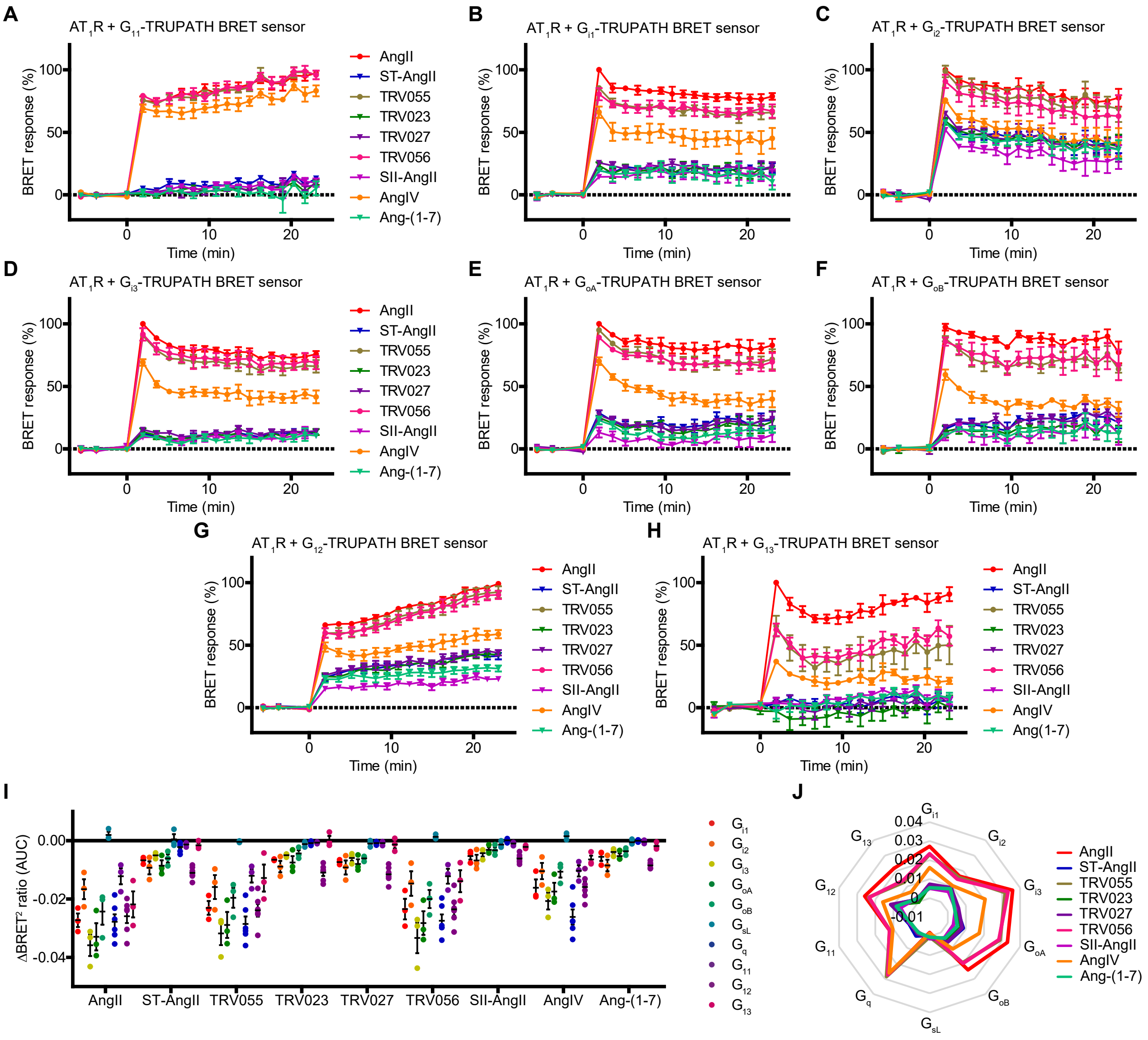


**Figure S3. Systematic assessment of the agonist-specific G protein activation profile of $AT_1R$ using the TRUPATH BRET sensor set**

**A–H** Activation kinetics of the indicated TRUPATH G protein BRET sensors. G protein activation is reflected by decrease of the BRET ratio. Positive values for BRET responses are achieved by expressing changes in the BRET ratio as a percentage of the highest effect induced by AngII. $AT_1R$ agonists were used at 10 µM.

**I–J** Scatter dot plot and radial plot show the average change in BRET ratio (AUC). Data are mean ± SEM in panels A–I, data are mean in panel J. $N = 6$ for $G_q$ and $G_{12}$ BRET sensors, $N = 3$ for the other G protein sensors.

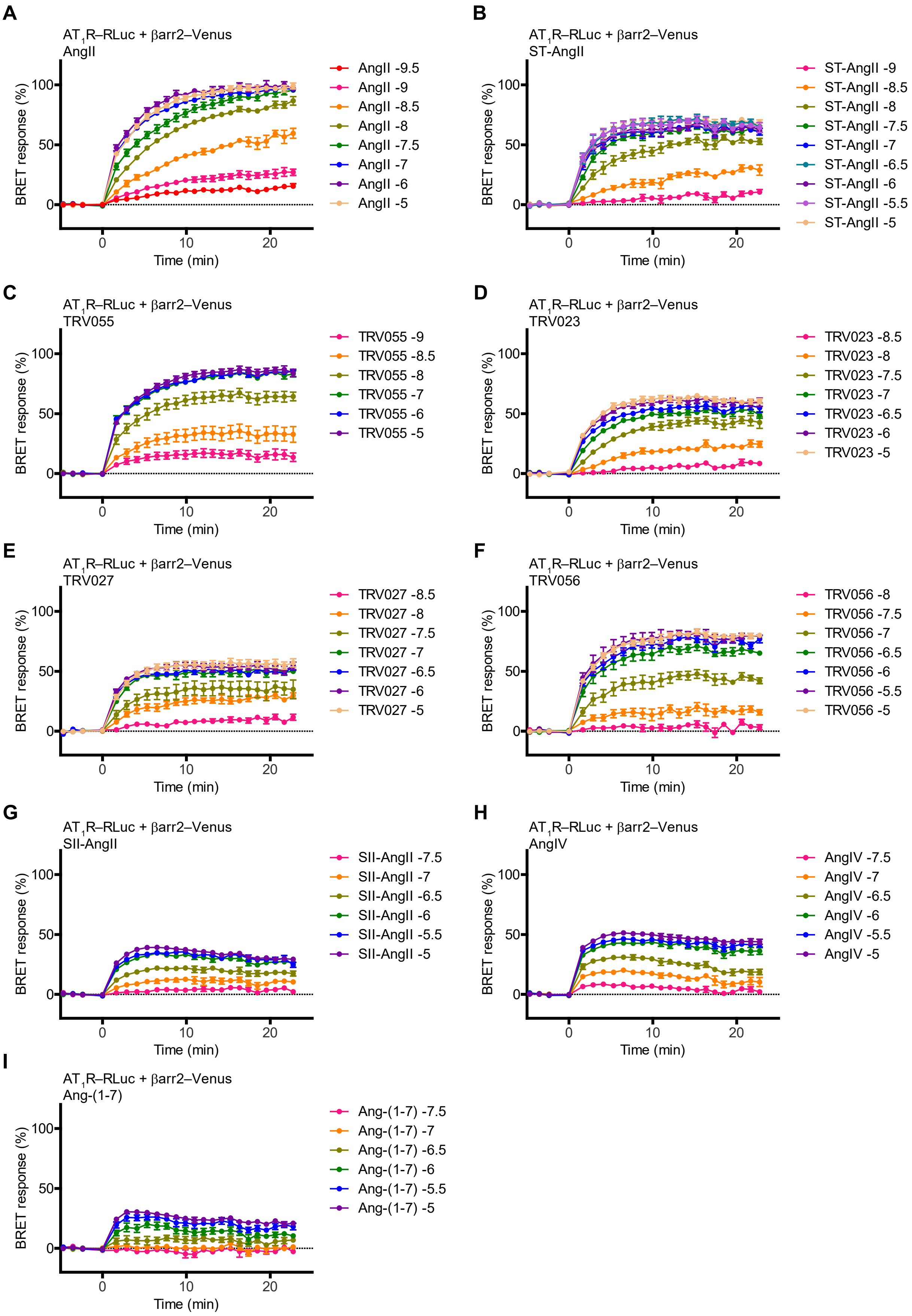
A
AT1R–RLuc + βarr2–Venus
AngII
AngII -9.5
AngII -9
AngII -8.5
AngII -8
AngII -7.5
AngII -7
AngII -6
AngII -5
B
AT1R–RLuc + βarr2–Venus
ST-AngII
ST-AngII -9
ST-AngII -8.5
ST-AngII -8
ST-AngII -7.5
ST-AngII -7
ST-AngII -6.5
ST-AngII -6
ST-AngII -5.5
ST-AngII -5
C
AT1R–RLuc + βarr2–Venus
TRV055
TRV055 -9
TRV055 -8.5
TRV055 -8
TRV055 -7
TRV055 -6
TRV055 -5
D
AT1R–RLuc + βarr2–Venus
TRV023
TRV023 -8.5
TRV023 -8
TRV023 -7.5
TRV023 -7
TRV023 -6.5
TRV023 -6
TRV023 -5
E
AT1R–RLuc + βarr2–Venus
TRV027
TRV027 -8.5
TRV027 -8
TRV027 -7.5
TRV027 -7
TRV027 -6.5
TRV027 -6
TRV027 -5
F
AT1R–RLuc + βarr2–Venus
TRV056
TRV056 -8
TRV056 -7.5
TRV056 -7
TRV056 -6.5
TRV056 -6
TRV056 -5.5
TRV056 -5
G
AT1R–RLuc + βarr2–Venus
SII-AngII
SII-AngII -7.5
SII-AngII -7
SII-AngII -6.5
SII-AngII -6
SII-AngII -5.5
SII-AngII -5
H
AT1R–RLuc + βarr2–Venus
AngIV
AngIV -7.5
AngIV -7
AngIV -6.5
AngIV -6
AngIV -5.5
AngIV -5
I
AT1R–RLuc + βarr2–Venus
Ang-(1-7)
Ang-(1-7) -7.5
Ang-(1-7) -7
Ang-(1-7) -6.5
Ang-(1-7) -6
Ang-(1-7) -5.5
Ang-(1-7) -5
BRET response (%)
Time (min)
0
50
100
10
20

**Figure S4. Kinetics of β-arrestin2–Venus binding to $AT_1R$–RLuc upon stimulation with increasing concentrations of agonists**

**A–I** Cells were stimulated with the indicated agonists. The numbers after the names of the ligands in the legends represent the log10 value of the agonist concentration. Data are mean ± SEM, *N* = 4–19.

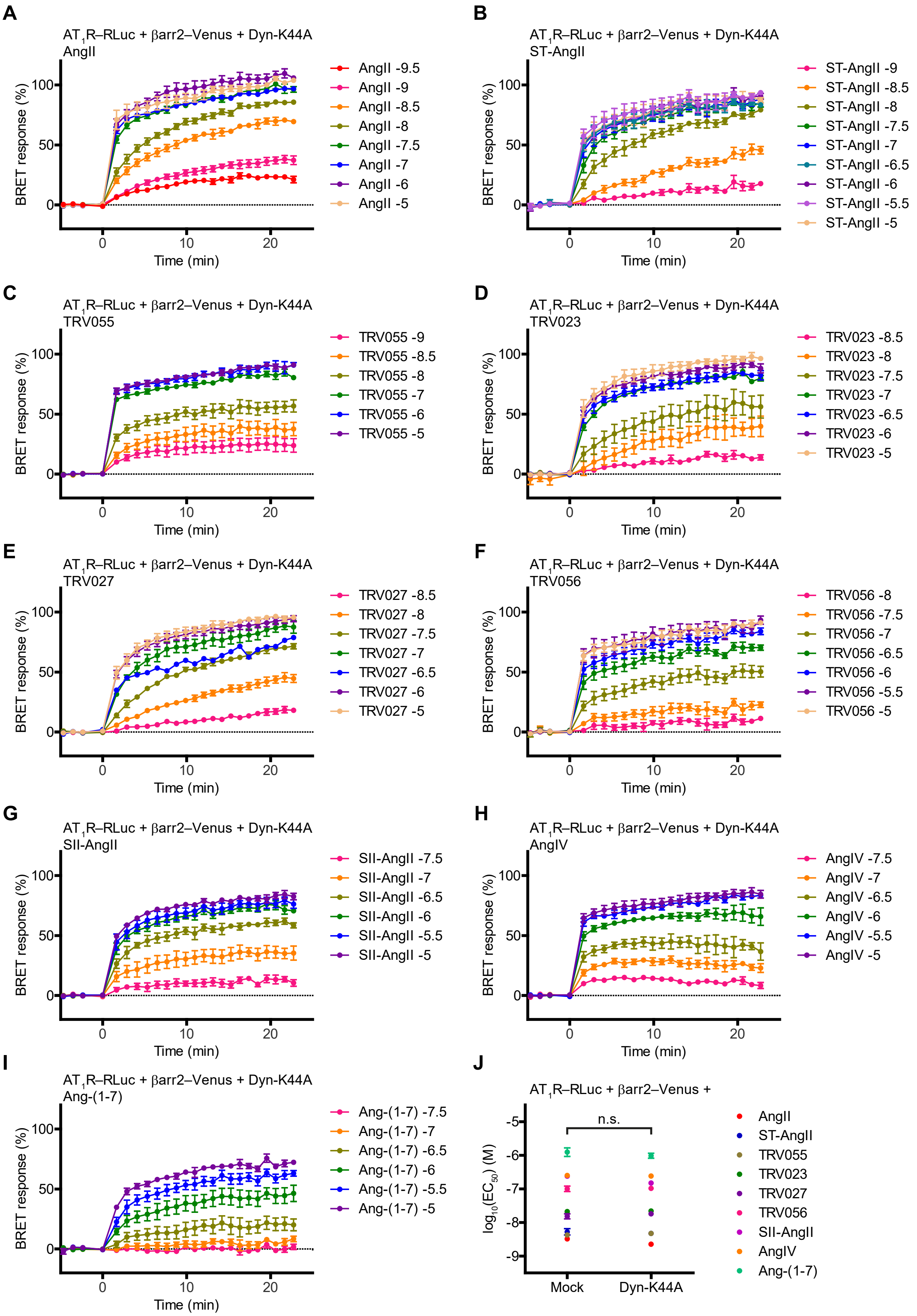
A
AT1R–RLuc + βarr2–Venus + Dyn-K44A
AngII
BRET response (%)
Time (min)
AngII -9.5
AngII -9
AngII -8.5
AngII -8
AngII -7.5
AngII -7
AngII -6
AngII -5
B
AT1R–RLuc + βarr2–Venus + Dyn-K44A
ST-AngII
BRET response (%)
Time (min)
ST-AngII -9
ST-AngII -8.5
ST-AngII -8
ST-AngII -7.5
ST-AngII -7
ST-AngII -6.5
ST-AngII -6
ST-AngII -5.5
ST-AngII -5
C
AT1R–RLuc + βarr2–Venus + Dyn-K44A
TRV055
BRET response (%)
Time (min)
TRV055 -9
TRV055 -8.5
TRV055 -8
TRV055 -7
TRV055 -6
TRV055 -5
D
AT1R–RLuc + βarr2–Venus + Dyn-K44A
TRV023
BRET response (%)
Time (min)
TRV023 -8.5
TRV023 -8
TRV023 -7.5
TRV023 -7
TRV023 -6.5
TRV023 -6
TRV023 -5
E
AT1R–RLuc + βarr2–Venus + Dyn-K44A
TRV027
BRET response (%)
Time (min)
TRV027 -8.5
TRV027 -8
TRV027 -7.5
TRV027 -7
TRV027 -6.5
TRV027 -6
TRV027 -5
F
AT1R–RLuc + βarr2–Venus + Dyn-K44A
TRV056
BRET response (%)
Time (min)
TRV056 -8
TRV056 -7.5
TRV056 -7
TRV056 -6.5
TRV056 -6
TRV056 -5.5
TRV056 -5
G
AT1R–RLuc + βarr2–Venus + Dyn-K44A
SII-AngII
BRET response (%)
Time (min)
SII-AngII -7.5
SII-AngII -7
SII-AngII -6.5
SII-AngII -6
SII-AngII -5.5
SII-AngII -5
H
AT1R–RLuc + βarr2–Venus + Dyn-K44A
AngIV
BRET response (%)
Time (min)
AngIV -7.5
AngIV -7
AngIV -6.5
AngIV -6
AngIV -5.5
AngIV -5
I
AT1R–RLuc + βarr2–Venus + Dyn-K44A
Ang-(1-7)
BRET response (%)
Time (min)
Ang-(1-7) -7.5
Ang-(1-7) -7
Ang-(1-7) -6.5
Ang-(1-7) -6
Ang-(1-7) -5.5
Ang-(1-7) -5
J
AT1R–RLuc + βarr2–Venus +
n.s.
log10(EC50) (M)
Mock
Dyn-K44A
AngII
ST-AngII
TRV055
TRV023
TRV027
TRV056
SII-AngII
AngIV
Ang-(1-7)

**Figure S5. Inhibition of endocytosis markedly changes the efficacy but not the potency of $AT_1R$ agonists in β-arrestin2 binding measurements**

**A–I** Kinetics of β-arrestin2–Venus binding to $AT_1R$–RLuc upon stimulation with increasing concentrations of agonists in Dyn-K44A coexpressing cells. The cells were stimulated with the indicated agonists, the numbers in the legends represent the log10 value of the agonist concentration. Data are mean ± SEM, $N = 3–16$.

**J** Comparison of log $EC_{50}$ values of β-arrestin2 recruitment in the absence or presence of DynK44A coexpression. The fitted log $EC_{50}$ values of Fig. 1E and F are shown. The means of agonist responses were compared with two-tailed paired t test, $P = 0.2199$.

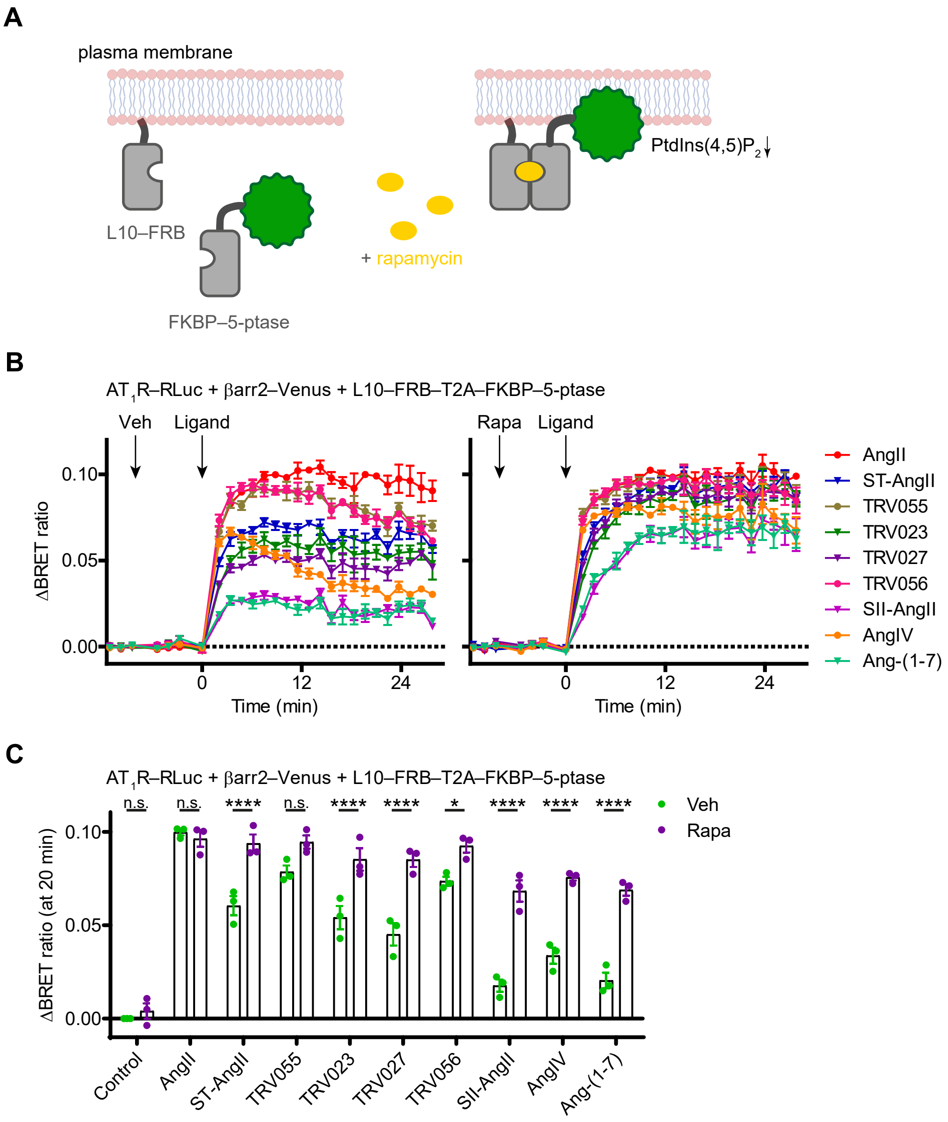


**Figure S6. Inhibition of receptor trafficking by PtdIns(4,5)$P_2$ depletion recapitulates the effects of endocytosis inhibition on $AT_1R$–β-arrestin2 interaction**

**A** Schematic representation of the PtdIns(4,5)$P_2$ depletion system. PtdIns(4,5)$P_2$ is an important regulator of endocytosis, and its depletion was shown to inhibit the clathrin-mediated receptor internalization. Therefore, cells were transfected with the L10–FRB–T2A–FKBP–5-ptase construct, from which L10–FRB (FRB targeted to the plasma membrane by the fusion tag L10) and FKBP–5-ptase (FKBP-fused 5-ptase) are translated in equimolar amounts due to the T2A sequence. Treatment with rapamycin (Rapa, 300 nM) induces heterodimerization between FRB and FKBP, thus FKBP–5-ptase is translocated to the plasma membrane where it can cleave PtdIns(4,5)$P_2$.

**B–C** PtdIns(4,5)$P_2$ depletion greatly reduced the ligand-specific differences in β-arrestin2 binding. In B, kinetic curves are shown, the arrows represent the time of pretreatment (vehicle (Veh) or Rapa) and agonist stimulation. Note that association rate was not reduced upon rapamycin treatment as it was in the case of hypertonic sucrose (Fig. S7C), suggesting that the different association kinetics observed with hypertonic sucrose is unrelated to endocytosis inhibition and sucrose may also exert direct effects on β-arrestin binding. Scatter dot plots of changes in BRET ratios at 20 min after stimulation are shown in C. Data are mean ± SEM, $N = 3$. Two-way ANOVA with Bonferroni post-hoc test was used for statistical analysis. As Control has non-normal distribution, ANOVA was performed with its exclusion. For AngII, $P > 0.9999$; for ST-AngII, TRV023, TRV027, SII-AngII, AngIV, and Ang-(1-7), ****, $P < 0.0001$; for TRV055, $P = 0.1171$; for TRV056, *, $P = 0.0327$. Rapamycin effect on control was analyzed with one-sample t test, whether mean differs from 0 (mean of Veh:Control condition), $P = 0.4402$.

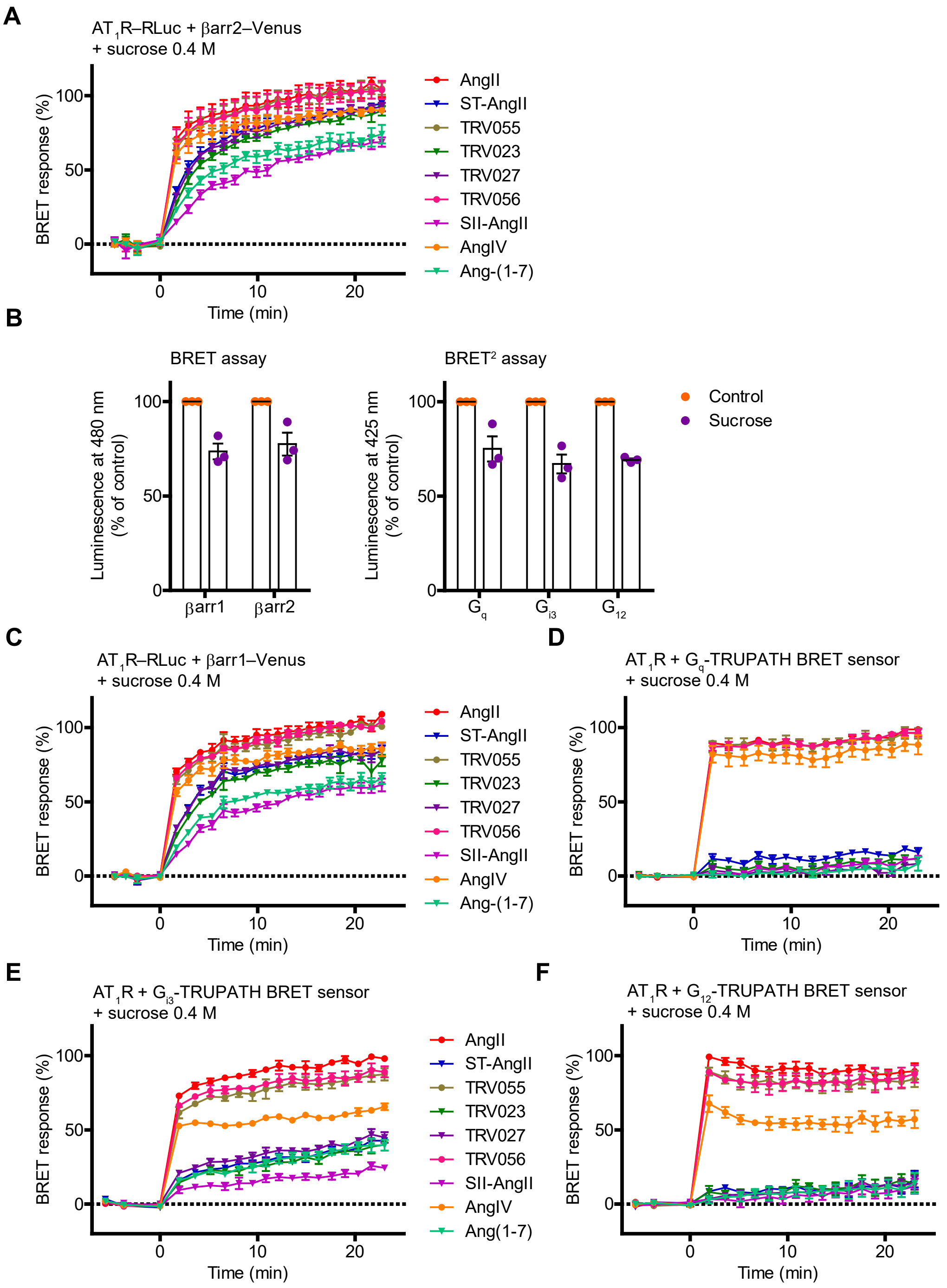


**Figure S7. Effect of endocytosis inhibition with hypertonic sucrose on the transducer activation profile of AT$_1$R agonists**

**A**, **C** Effect of hypertonic sucrose on the kinetics of β-arrestin2–Venus (A) and β-arrestin1-Venus (C) binding to AT$_1$R–RLuc. The control curves are shown in Fig. 1A and Fig. S2A. Notably, we also found changes in the β-arrestin association kinetics, but this effect of hypertonic sucrose is probably unrelated to receptor internalization.

**B** Hypertonic sucrose (0.4 M) decreased the luminescence values at both 480 nm and 530 nm wavelengths, which can alter the basal signals and the changes in the BRET ratio. Therefore, agonist responses were compared only under the same experimental conditions (i.e. with or without sucrose). Pooled analysis of luminescence intensity values for BRET and $BRET^2$ setups showed that sucrose treatment significantly decreased the luminescence intensities (one-sample test, mean significantly differs from 100, ***, P = 0.0008 for BRET and ****, P <0.0001 for $BRET^2$.)
**D–F** Kinetics of activation of $G_q$, $G_{i3}$ and $G_{12}$ protein BRET sensors in the presence of hypertonic sucrose. The control curves are shown in Fig. 1B, S3D, and S3G.
Data are mean ± SEM, *N* = 3.

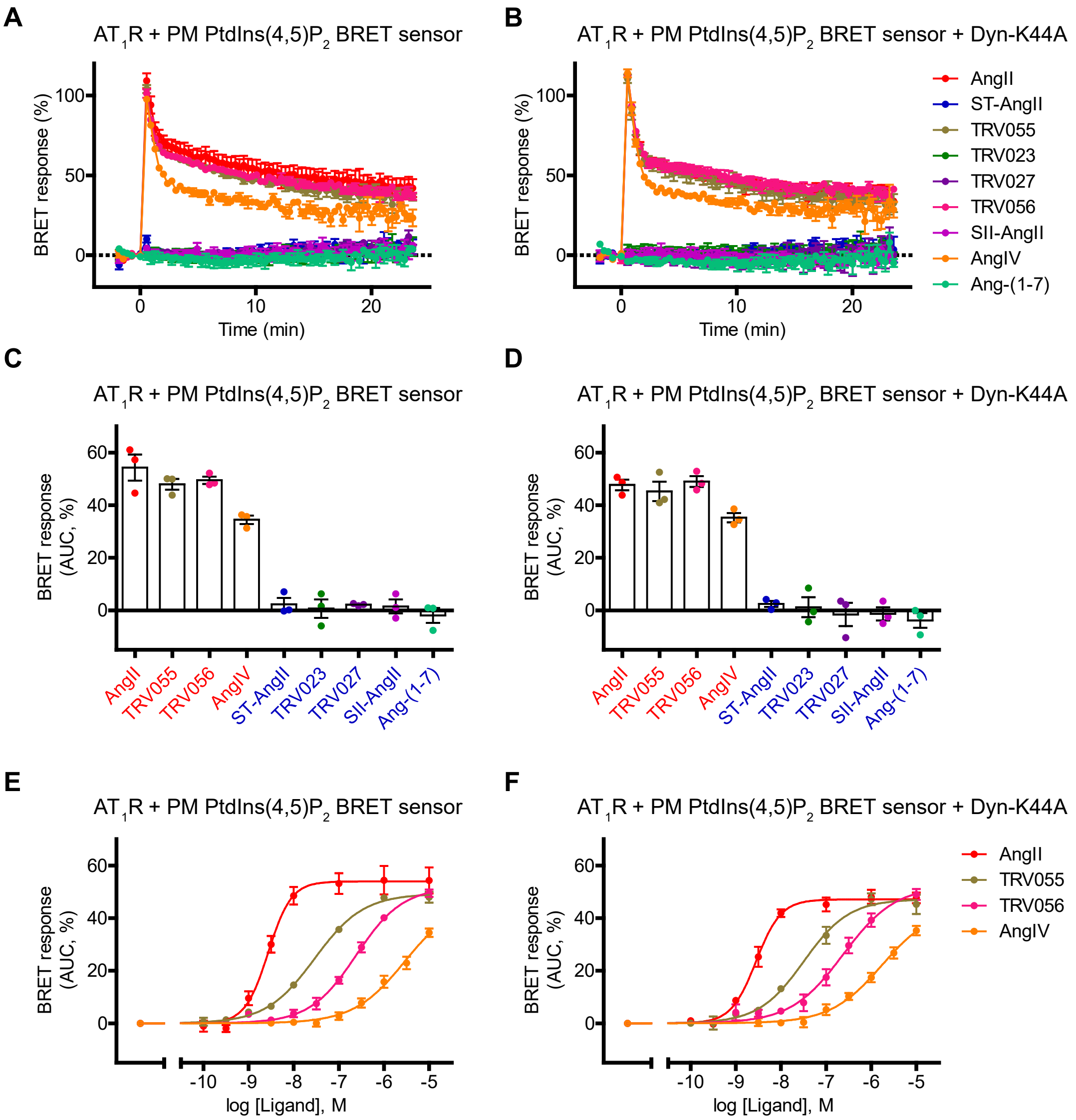


**Figure S8. $G_{q/11}$–PLCβ signaling pathway is not significantly affected by the blockade of endocytic processes**

PLCβ-mediated PtdIns(4,5)$P_2$ cleavage was monitored using a plasma membrane PtdIns(4,5)$P_2$ BRET sensor, where a decrease of PtdIns(4,5)$P_2$ is reflected by a decline in the BRET ratio. BRET responses were expressed as a percentage of the peak change in BRET ratio induced by 100 nM AngII (curve not shown) in each condition and were therefore positive values.

**A**–**B** Kinetic changes in plasma membrane PtdIns(4,5)$P_2$ levels with or without Dyn-K44A coexpression. All agonists were applied at 10 µM concentrations.

**C**–**D** AUC values from curves in panels A–B. Only the $G_q$-cluster ligands induced substantial PtdIns(4,5)$P_2$ cleavage and no significant effect of Dyn-K44 coexpression was found (two-way ANOVA with Bonferroni post-hoc test was perfomed, effect of Dyn-K44A coexpression was not significant ($P = 0.1685$), whereas stimulus had significant effect ****, $P <0.0001$), and no interaction was found ($P=0.9292$).

**E**–**F** Concentration–response curves of the $G_q$-activating ligands in cells expressing the indicated constructs.

Data are mean ± SEM, $N = 3$.

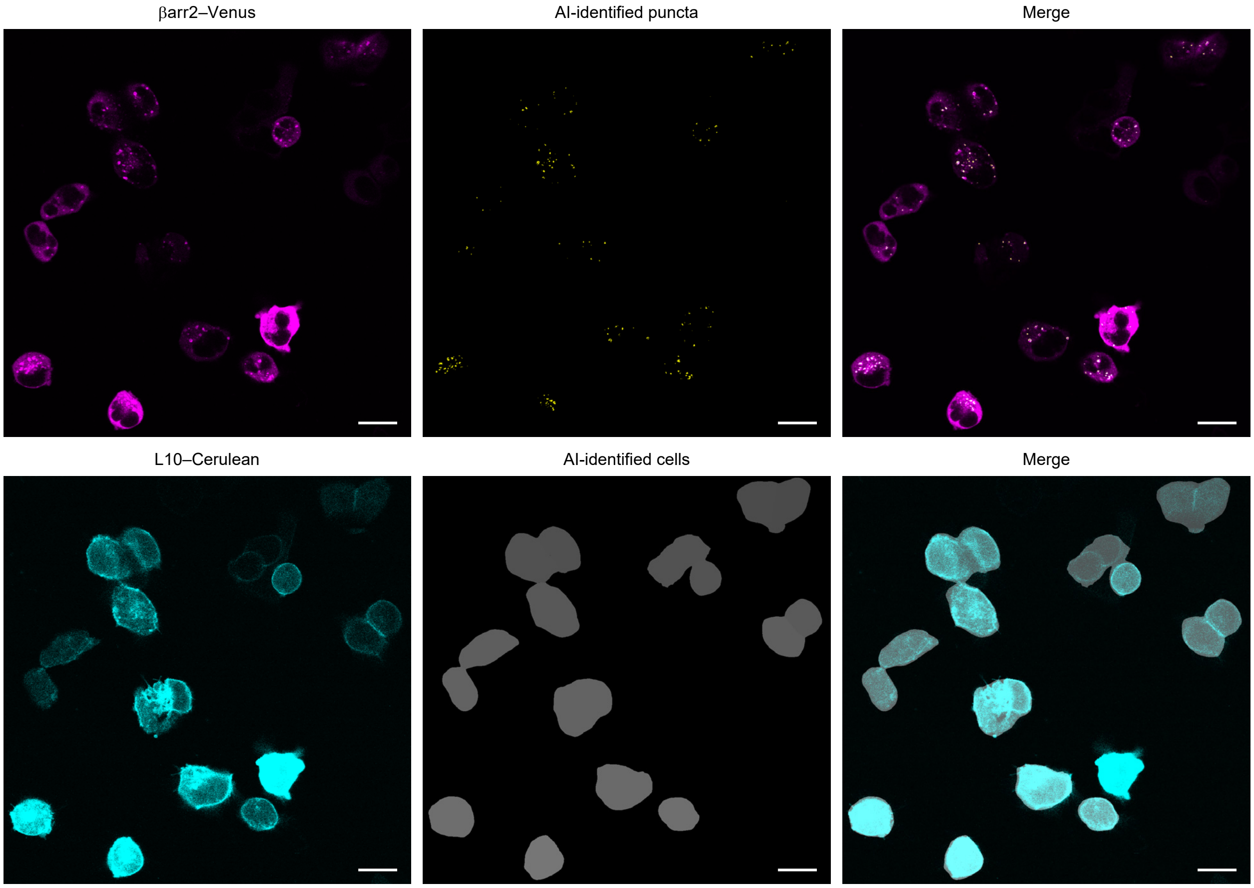


**Figure S9. Unsupervised, machine learning-based method for the detection of endosomal translocation of β-arrestin2–Venus**

Representative confocal images of live HEK 293A cells co-expressing $AT_1R$, β-arrestin2–Venus, and plasma membrane-targeted Cerulean (L10–Cerulean). Images were acquired 20–30 min after stimulation with 10 µM AngII. L10–Cerulean was used to mark the contour of cells. Top images display the intracellular β-arrestin2-Venus fluorescence and the puncta identified by the neural network-based algorithm. Bottom images show how cells were detected based on the fluorescent plasma membrane marker by the Cellpose algorithm. Scale bars are 20 µm. Note, that in contrast to live cell images, puncta formation could be observed in control, unstimulated fixed cells as well, suggesting that fixation may cause artificial aggregations.

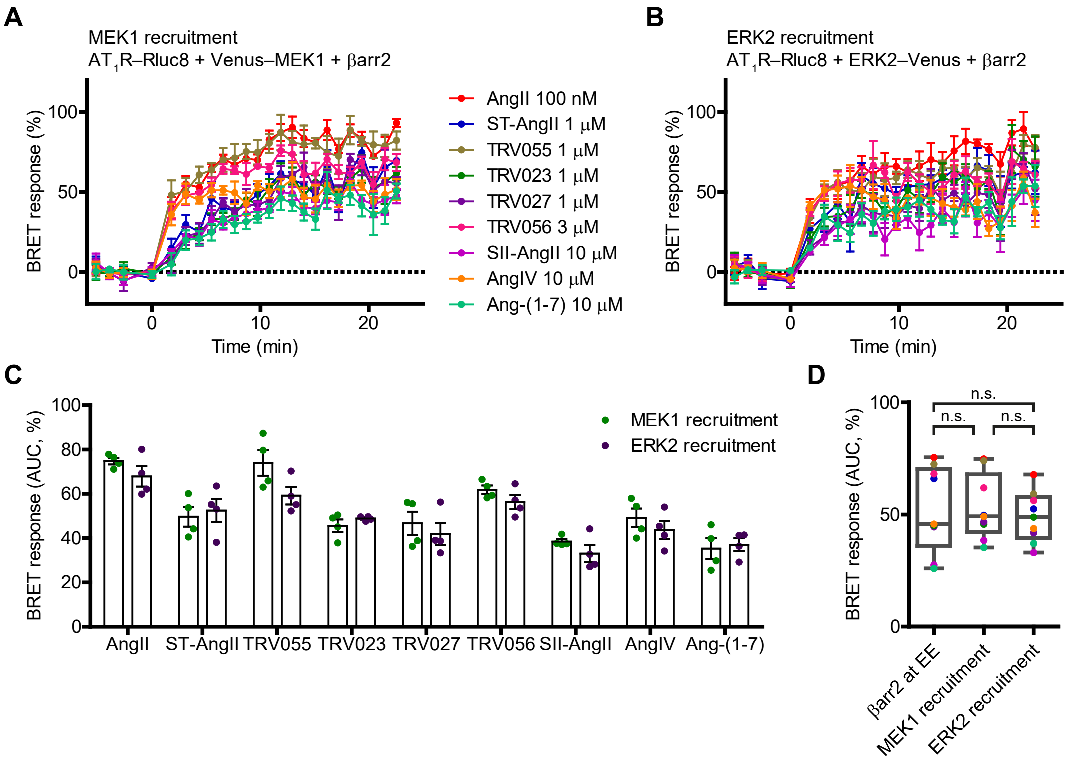


**Figure S10. Real-time monitoring of $AT_1R$–β-arrestin2–MEK1/ERK2 complex formation with BRET**

**A–B** To follow the MEK1 or ERK2 recruitment, HEK 293T cells were co-transfected with the indicated constructs. The agonists were used at the indicated concentrations.

**C** Scatter dot plot shows the AUC values of the curves from A and B. $N = 4$.

**D** Distribution of the signals of the ligands in the indicated BRET assays, the corresponding kinetic curves are shown in Fig. 2C, S10A and S10B. F tests were performed on average normalized datasets, βarr2 at EE vs. MEK1 recruitment, P = 0.4345; βarr2 at EE vs. ERK2 recruitment, P = 0.2177; MEK1 recruitment vs. ERK2 recruitment, P = 0.6419.

Data are mean ± SEM in A–C, and data are mean in D.

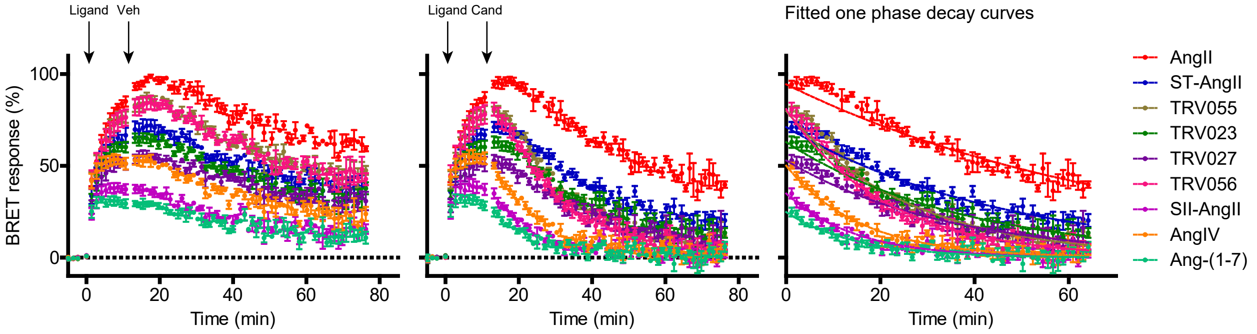


**Figure S11. Monitoring the β-arrestin2 dissociation from $AT_1R$ after agonist displacement** BRET was continuously monitored in cells expressing $AT_1R$–Rluc and β-arrestin2–Venus. After agonist stimulation (first arrow), displacement of agonists was induced by candesartan (Cand, 10 µM) treatment (second arrow). Agonist concentrations were ~$EC_{50}$ × 30 (99 nM AngII, 145 nM ST-AngII, 148 nM TRV055, 511 nM TRV023, 436 nM TRV027, 2.6 µM TRV056, 7.31 µM SII-AngII, 7.25 µM AngIV, 25 µM Ang-(1-7)). Vehicle control is shown in the left panel. Right panel: one phase decay curves were fitted on the data points after candesartan treatment (time point 0) to assess $k_{dis}$ values for each ligand. Data are mean ± SEM, $N = 3$.

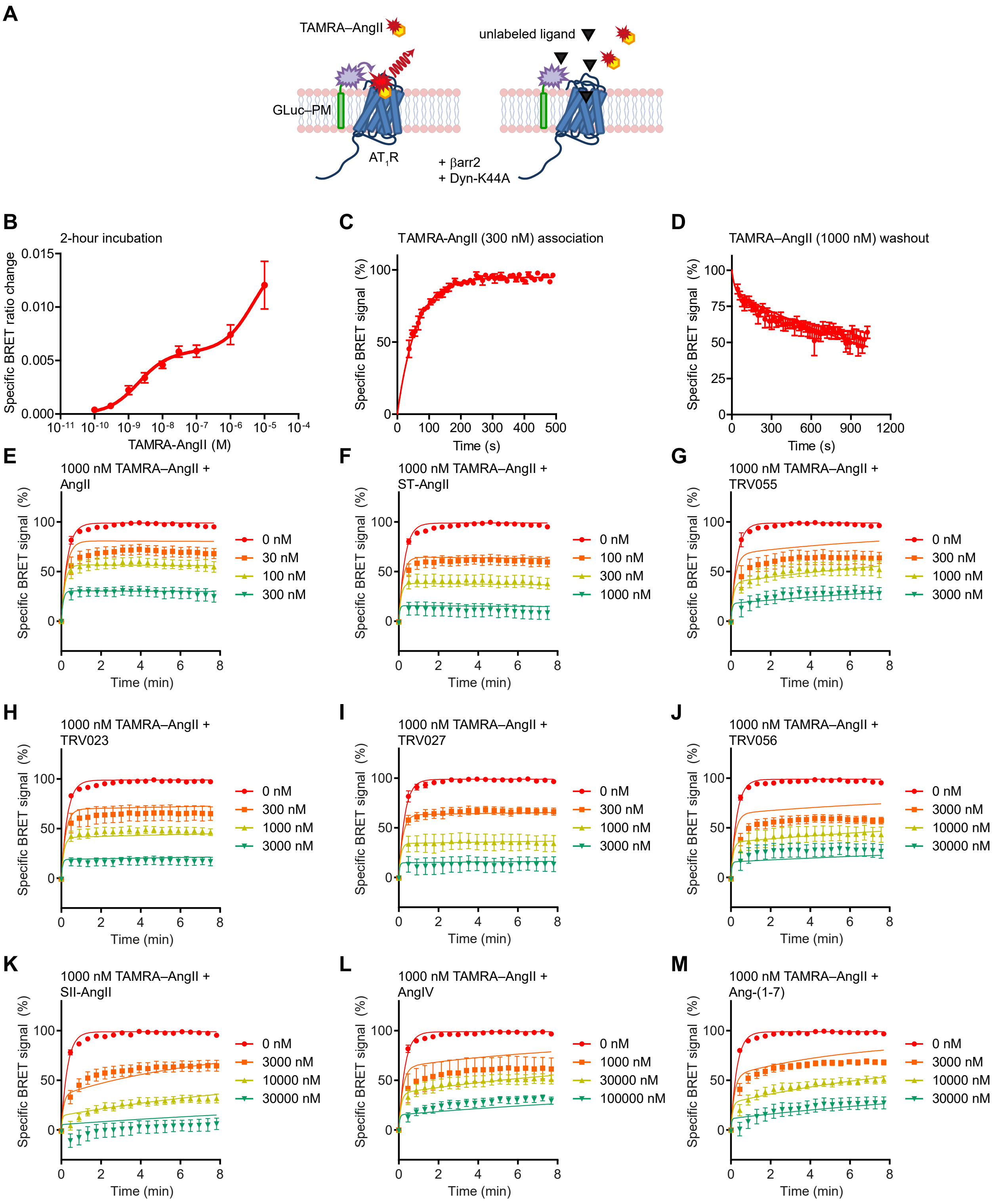


**Figure S12. BRET-based measurement of the kinetic rate constants of ligand–receptor interaction in cells overexpressing β-arrestin2**

**A** Schematic representation of the BRET-based competitive binding assay for kinetic ligand binding measurements. Cells were co-transfected with $AT_1R$, β-arrestin2, Dyn-K44A, and cell

surface-targeted *Gaussia* luciferase (GLuc–PM) biosensor (*33*). Upon binding of TAMRA-labeled AngII (TAMRA–AngII) to $AT_1R$, the enrichment of the fluorophore at the plasma membrane leads to a specific increase of the bystander BRET signal. This signal is reduced if the same binding site is occupied by a non-labeled orthosteric $AT_1R$ ligand.
**B** Specific change in BRET ratio upon increasing concentration of TAMRA–AngII treatment after 2-h incubation at room temperature. Two-site specific binding curve was fitted ($B_{max\_high}$ = 0.0058, $K_{d\_high}$ = 1.96 nM, $B_{max\_low}$ = 0.0091, $K_{d\_low}$ = 4.68 μM).
**C**–**D** Monitoring the association (C) and dissociation (D) kinetics between TAMRA–AngII and $AT_1R$. TAMRA–AngII was used in 300 nM in panel C. This concentration allows to capture ample measurement points before reaching the steady state. To assess the dissociation kinetics, 1 μM TAMRA–AngII treatment was applied for 15 min and the BRET signal was monitored after washout of TAMRA–AngII. 10 μM candesartan was added to prevent rebinding of TAMRA–AngII. In D, dissociation – 2 binding site equation was used ($k_{off\_high} = 3.45 \times 10^{-2}$ $min^{-1}$, $k_{off\_low}$ = 2.78 $min^{-1}$). Since at the applied TAMRA–AngII concentration mostly the high-affinity binding site is occupied, $k_{off\_high}$ was applied as $k_{off\_LR}$ for TAMRA–AngII in the further calculations, and one-site binding models were used for simplification. Association kinetics (one conc. of hot) equation was applied to calculate kon_LR for TAMRA–AngII ($k_{on\_LR} = 3.24 \times 10^6$ $M^{-1}$ $min^{-1}$).
**E**–**M** Competitive kinetic ligand binding measurements to assess $k_{on\_LR}$ and $k_{off\_LR}$ values for the non-labeled $AT_1R$ ligands. Cells were simultaneously treated with 1 μM TAMRA–AngII and the indicated ligands at the indicated concentrations. Kinetics of competitive binding equations were fitted. The fitted $k_{on\_LR}$ and $k_{off\_LR}$ values for the non-labeled ligands are shown in Table 1.
In all panels, non-specific signal was assessed by cotreatment with 10 μM candesartan and was subtracted from the total signal. Data are mean ± SEM, *N* = 3–5.

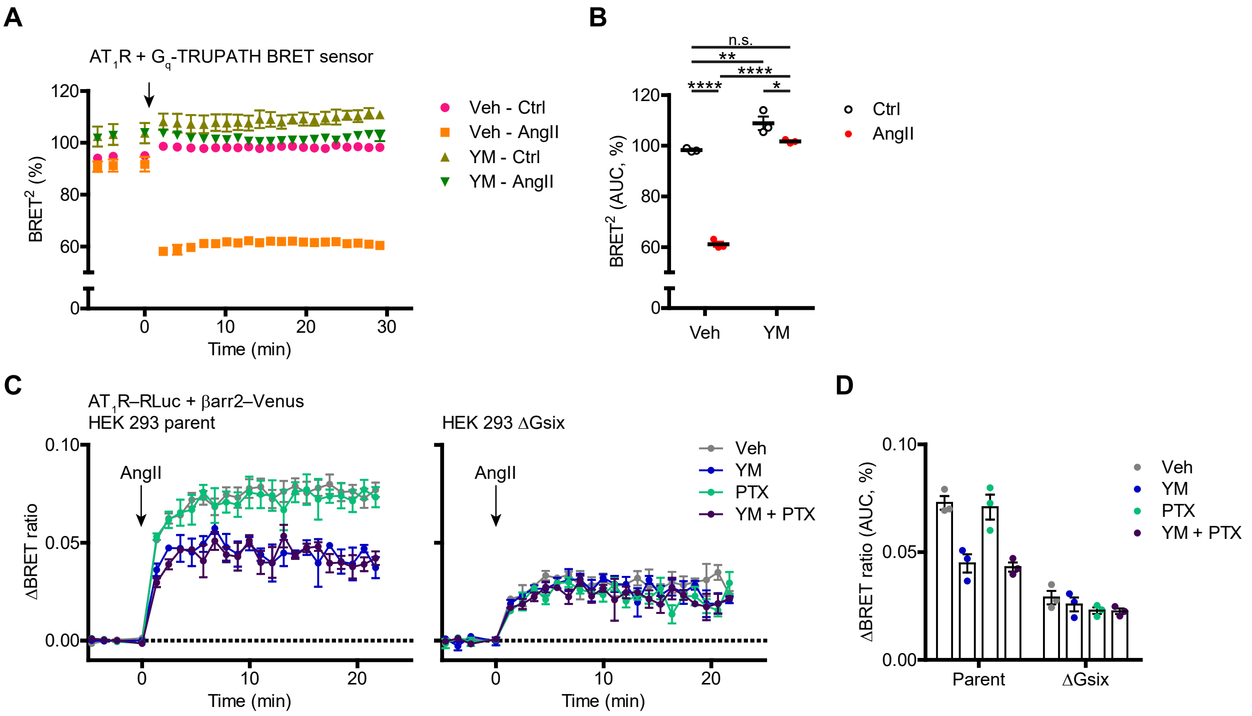


**Figure S13. $G_{q/11}$-inhibitor YM-254890 effectively and selectively reduced AngII-induced $G_q$ protein activity**

**A–B** Cells coexpressing $AT_1R$ and the $G_q$ TRUPATH sensor subunits were pretreated with 100 nM YM-254890 for 40 minutes. YM pretreatment inhibited the AngII (10 µM)-induced activation of the $G_q$ TRUPATH BRET sensor and reduced its baseline activity. $BRET^2$ ratios are expressed as the percentage of the highest measured ratio in vehicle-pretreated control cells in each experiment.

**C–D** AT1R–Rluc–β-arrestin2–Venus interaction was monitored in parent and ΔGsix HEK 293 cells, 40-min YM (100 nM) and/or 20-h PTX (100 ng/ml) pretreatments were applied for $G_{q/11}$ and $G_{i/o}$ protein inhibition, respectively. 10 µM AngII was used as stimuli. YM effects were absent in the G protein knockout cell line.

Kinetic curves are shown in A and C, AUC values of the corresponding curves are shown in B and D. $N = 3$, data are mean ± SEM.

In B, two-way ANOVA with Bonferroni post-hoc test was applied for statistical evaluation. Sources of variation: stimulus, pretreatment, and interaction, ****, $P < 0.0001$. Results of multiple comparisons: Veh:Control vs. Veh:AngII, Veh:AngII vs. YM:Control, and Veh:AngII vs. YM:AngII, ****, $P < 0.0001$; Veh:Control vs. YM:Control, **, $P = 0.0049$; Veh:Control vs. YM:AngII, n.s., $P = 0.7728$; YM:Control vs YM:AngII, *, $P = 0.048$.

Data of D was analyzed using three-way ANOVA. Sources of variation: cell type, ****, $P < 0.0001$; PTX, n.s, $P = 0.1944$; YM, ****, $P < 0.0001$. Interactions: cell type – PTX, n.s., $P = 0.5727$; cell type – YM, ****, $P < 0.0001$; PTX – YM, n.s., $P = 0.7485$; cell type – PTX – YM, n.s., $P = 0.7904$.

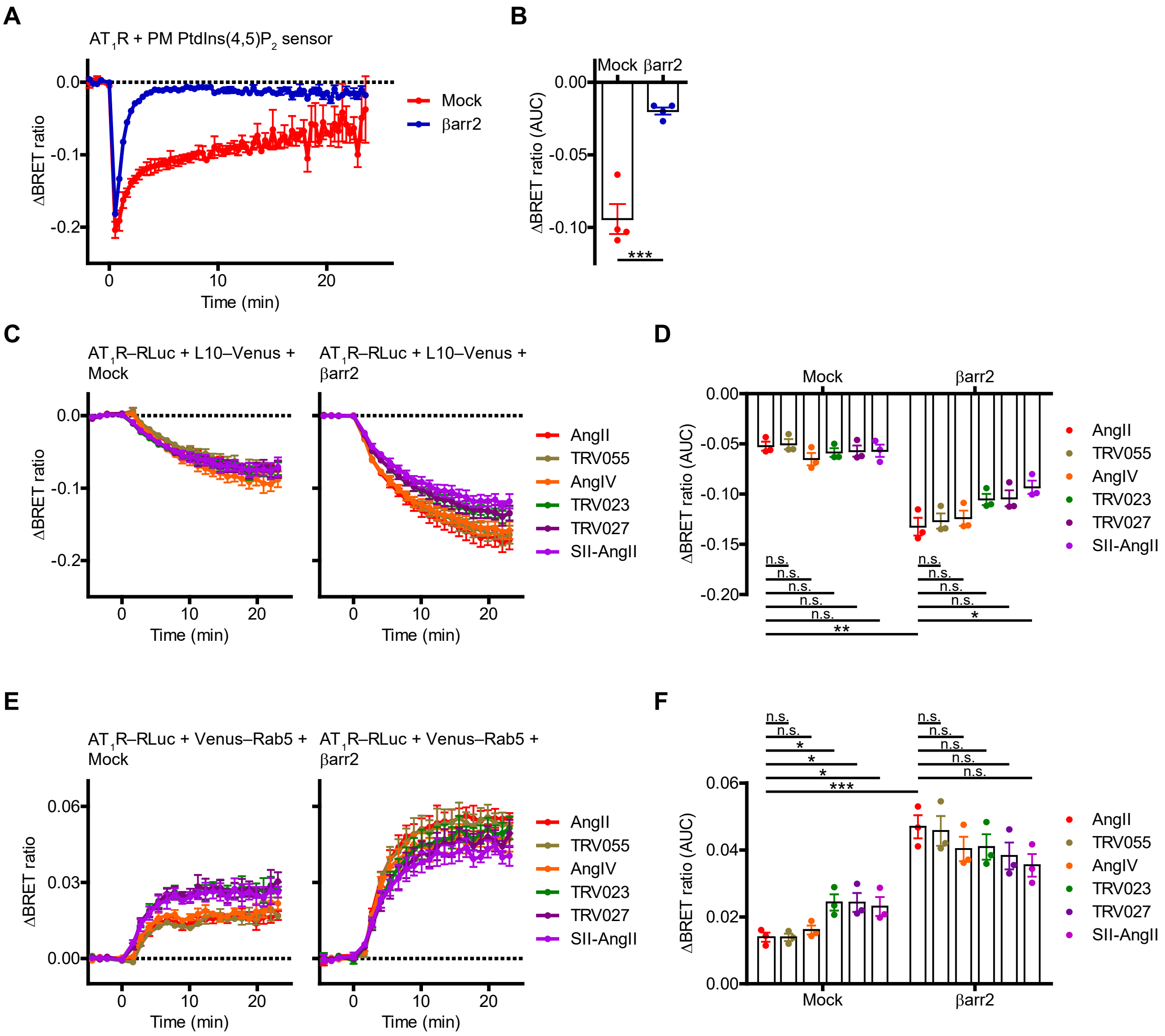


**Figure S14. Lack of PtdIns(4,5)$P_2$ cleavage-mediated inhibition of endocytosis in β-arrestin2 overexpressing cells**

**A–B** Decrease of the plasma membrane PtdIns(4,5)$P_2$ level after stimulation with 100 nM AngII, in cells cotransfected with $AT_1R$, plasma membrane PtdIns(4,5)$P_2$ BRET sensor, and empty pcDNA3.1 (Mock) or untagged β-arrestin2. In β-arrestin2-coexpressing cells, the PtdIns(4,5)$P_2$ decrease was more transient, reflecting an increased desensitization of the receptor. Kinetic curves are shown in A. Statistical analysis was performed on the AUC values of curves using unpaired two-tailed t test: ***, P = 0.0004 (B). $N = 4$.

**C–F** $AT_1R$ internalization was measured using bystander BRET assays in cells coexpressing the indicated constructs. In C–D, BRET was monitored between $AT_1R$–RLuc and the plasma membrane marker L10–Venus, where the decrease of BRET ratio indicates the translocation of $AT_1R$–RLuc from the plasma membrane to intracellular compartments. In E–F, BRET was followed between $AT_1R$–RLuc and the early endosome marker Venus–Rab5. In this assay, the increase of the BRET ratio indicates the translocation of $AT_1R$–RLuc to the Venus–Rab5-enriched early endosomes. The applied agonists were AngII (100 nM), TRV055 (1 μM), AngIV (10 μM),

TRV023 (1 μM), TRV027 (1 μM), and SII-AngII (10 μM). C and E show the kinetic curves, D and F represent the average changes in BRET ratios (AUC values), $N = 3$.

Earlier we showed that $G_{q/11}$ activation delays the endocytosis of $AT_1R$ because of the $G_{q/11}$ activity-related decrease of plasma membrane PtdIns(4,5)$P_2$, an important mediator of internalization (*29*). However, in β-arrestin2-coexpressing cells, in which the PtdIns(4,5)$P_2$ decrease is only transient, the $G_q$-cluster peptides did not induce less $AT_1R$ internalization, contradicting that their higher β-arrestin2 binding is due to suppressed endocytosis. In addition, only minor agonist related-differences in $AT_1R$ trafficking were observed in cells coexpressing β-arrestin2. Agonists were compared with the reference ligand AngII in each experimental conditions using one-way ANOVA with Bonferroni post-hoc test. AngII effects in Mock and β-arrestin2-coexpressing cells were compared with unpaired, two-tailed t-tests.

Data are mean ± SEM, n.s.: $P > 0.05$; *, $0.05 \geq P > 0.01$; **, $0.01 \geq P > 0.001$; ***, $0.001 \geq P > 0.0001$, ****, $P \leq 0.0001$.

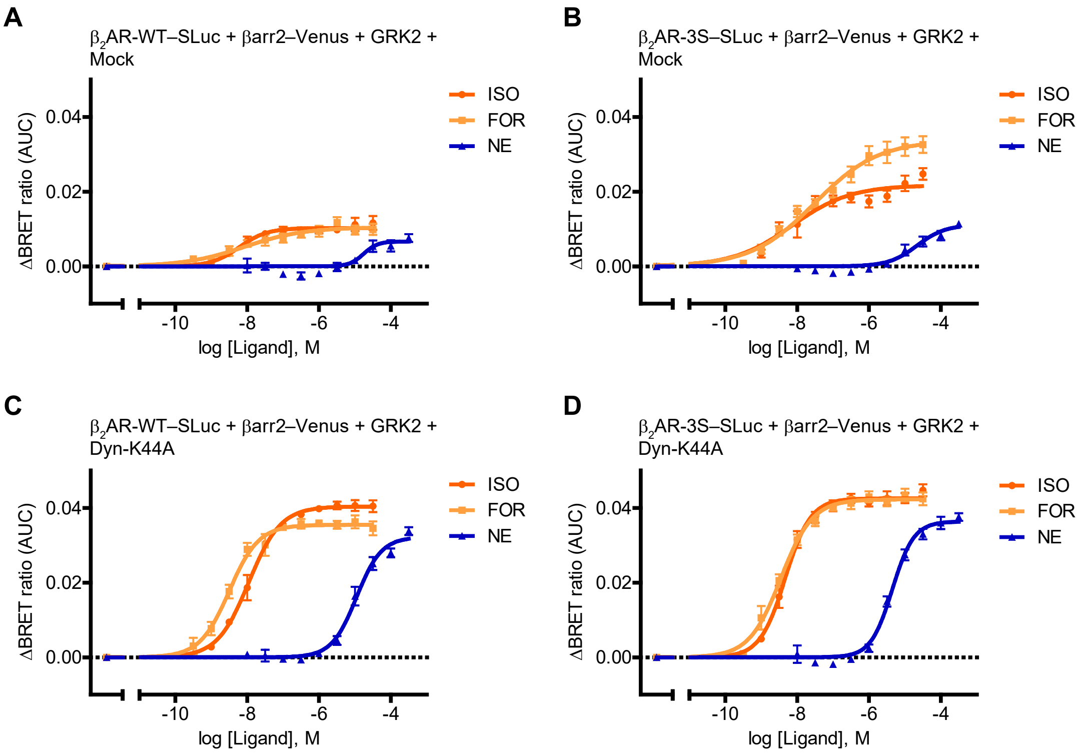


**Figure S15. Agonist- and endocytosis-dependent differences in β-arrestin2 recruitment of a β2AR mutant with engineered phosphorylation sites**

**A**–**D** Cells were cotransfected with the indicated constructs. BRET was measured upon stimulation with increasing concentrations of agonists (formoterol (FOR), isoproterenol (ISO), norepinephrine (NE)). Average changes in BRET ratios (AUC values) are shown as a function of ligand concentration. Corresponding kinetic curves of maximal ligand concentrations are shown in Fig. 5. Data are mean ± SEM, log(agonist) vs. response – variable slope curves were fitted. $N = 3$ for β2AR–SLuc, $N = 4$ for β2AR-3S–SLuc measurements.

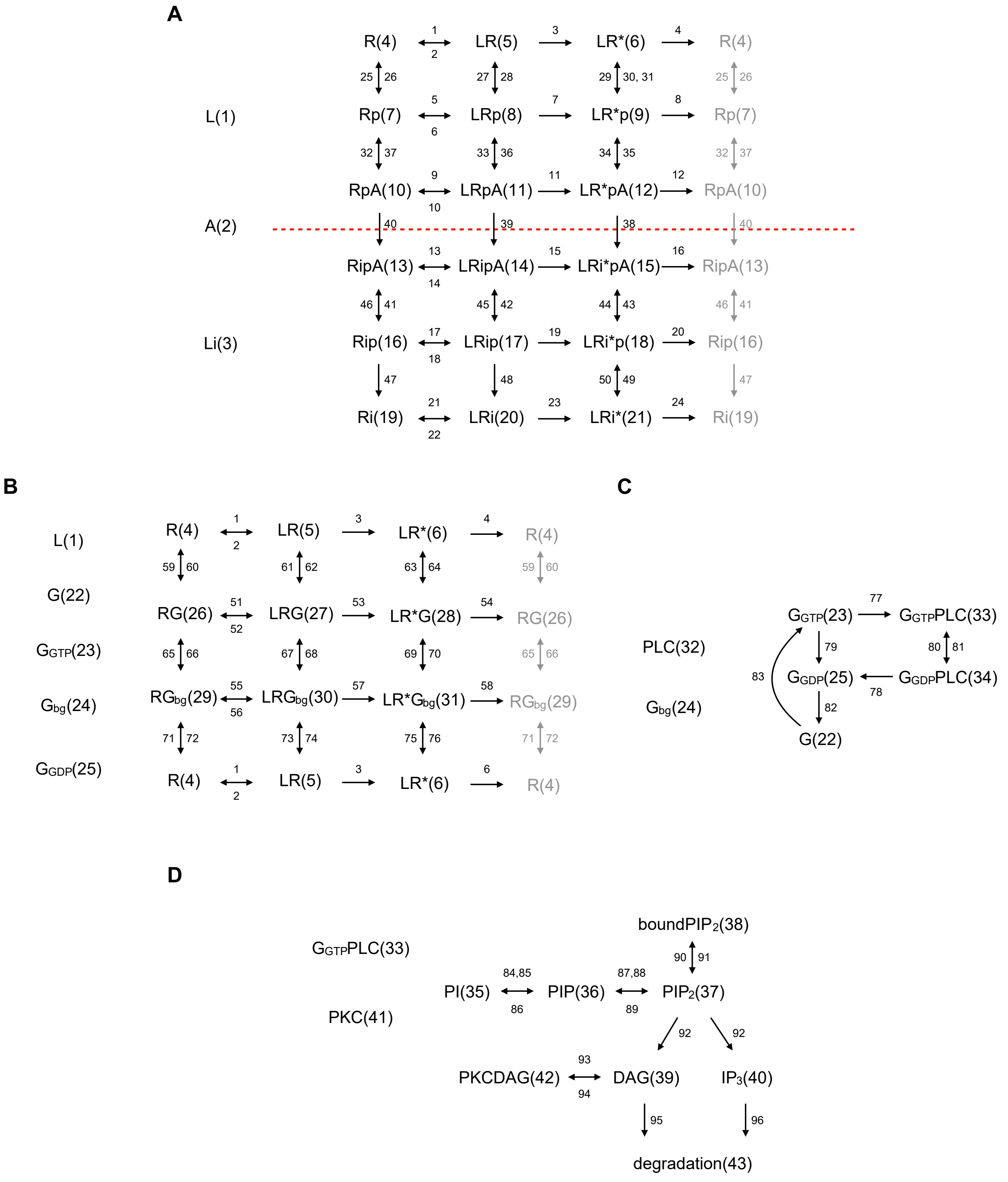


**Figure S16. Schematic representation of the molecules and reactions incorporated into the ODE model**

Numbers in parentheses indicate the index of molecules. Double-headed arrows depict reversible reactions, where the index of the rate constant of the reaction from left to right is indicated above the arrow, and the number below the arrow denotes the index of the rate constant of the complementary reaction. Single-headed arrows denote irreversible reactions, and the index of their corresponding rate constants are indicated above the arrows.

**A** β-arrestin pathway

**B** G protein activation

**C** and **D** Phospholipase C activation and second messenger generation

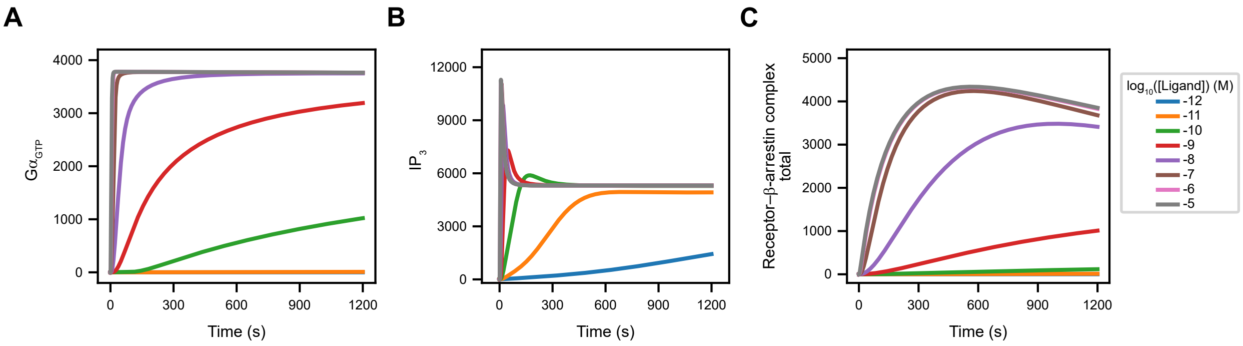


**Figure S17. Simulated time course profiles of major GPCR signaling events**

Simulated time courses of G protein activation (**A**, $G_{GTP}$ refers to molecules 23, 33), $IP_3$ formation (**B**, $IP_3$: molecule 40), and total receptor–β-arrestin binding (**C**, molecules 10, 11, 12, 13, 14, 15) upon stimulation with increasing concentration of a test agonist. Simulations were performed with a "low $k_{off}$" agonist ($k_{off\_LR}$ = 0.0003), legends indicate the $\log_{10}$ (agonist) concentrations in M, with the same color code applied across all panels. As in Methods thoroughly described, G protein activation was simulated in a system with overexpressed receptors & G proteins, $IP_3$ formation was modeled in a system with receptor overexpression, and β-arrestin binding was assessed in a system with receptor & β-arrestin overexpression.

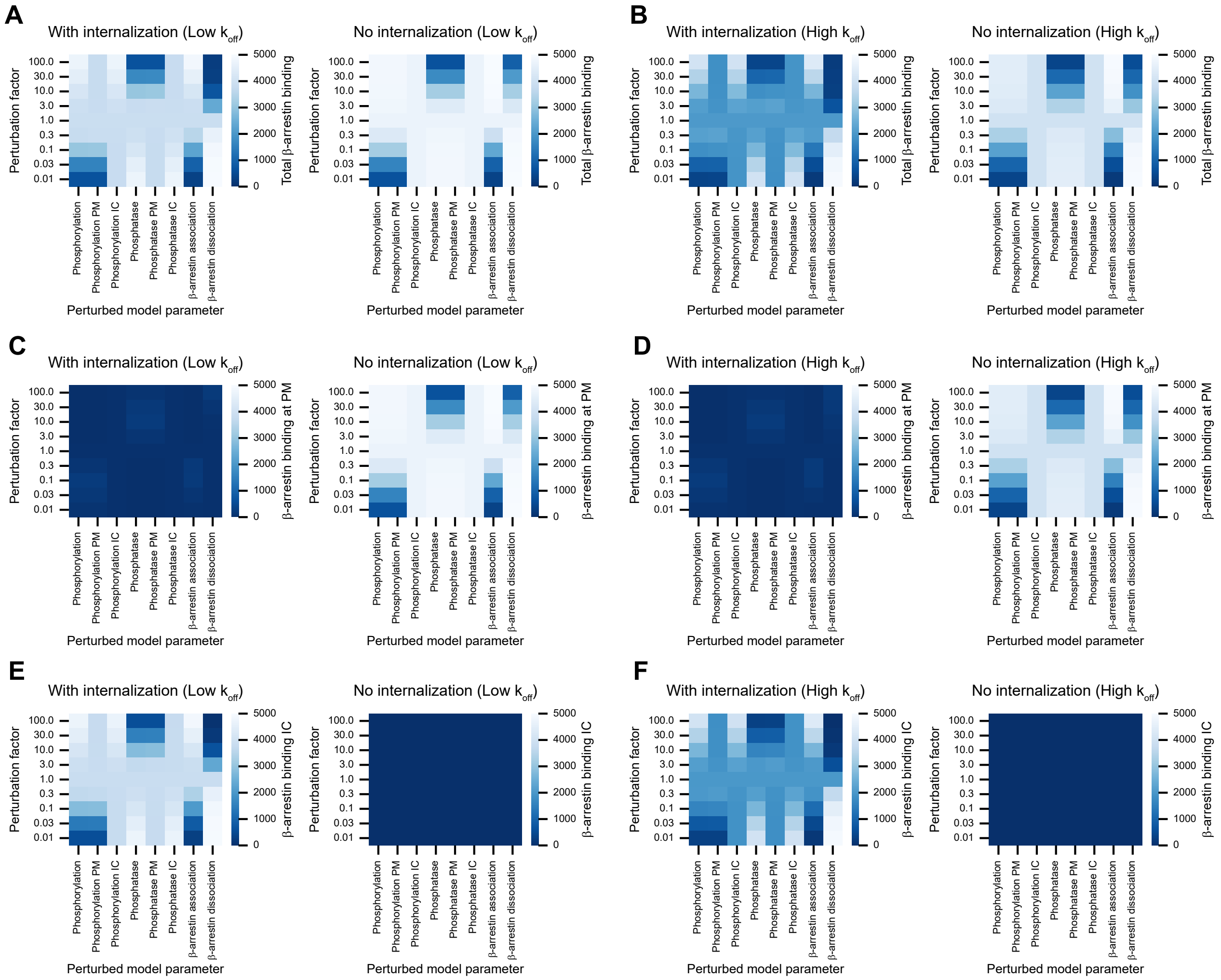


**Figure S18. Effects of perturbation of reaction rate constants of the β-arrestin binding pathway**

Changes in total (**A**, **B**), plasmalemmal (**C**, **D**) and endosomal (**E**, **F**) β-arrestin binding after perturbation of reaction rate constants related to the β-arrestin binding pathway. Simulations were performed with a low $k_{off}$ ($k_{off\_LR} = 0.0003$; **A**, **C**, **E**) and a high $k_{off}$ ($k_{off\_LR} = 0.03$; **B**, **D**, **F**) agonist. The perturbated factors are described in the Methods section and briefly in the legend of Fig. 6E. Left panels show the results of simulations with permitted internalization, the right panels present the results without internalization.

**Table S1. Initial concentration values of modeled molecular species.** Table includes molecule index (from Fig. S16) and initial concentration (with unit) for each molecular species. Comments describe rationale for selecting initial concentration in case of absence / discrepancy from literature reference.

| | Molecule | Compartment | Description | Initial value | Unit | Comment | Reference |
|---|---|---|---|---|---|---|---|
| 1 | L | plasma membrane | Ligand | 0.0 | μM | constant during a single simulation | |
| 2 | A | plasma membrane | Arrestin | 15000.0 | molecule/$\mu m^2$ | corresponds to b-arrestin overexpression system (15 μM), converted to molecule/$\mu m^2$, assuming that cell volumen is 2,500 $\mu m^3$ and cell surface is 1,500 $\mu m^2$ | Ref. #41 |
| 3 | Li | endosomes | Ligand | 0.0 | μM | equals L(1) | |
| 4 | R | plasma membrane | Receptor | 5000.0 | molecule/$\mu m^2$ | corresponds to receptor overexpression system | Ref. #41 |
| 5 | LR | plasma membrane | Ligand-bound receptor | 0.0 | molecule/$\mu m^2$ | | |
| 6 | LR* | plasma membrane | Ligand-bound, active receptor | 0.0 | molecule/$\mu m^2$ | | |
| 7 | Rp | plasma membrane | Phosphorylated receptor | 0.0 | molecule/$\mu m^2$ | | |
| 8 | LRp | plasma membrane | Ligand-bound, phosphorylated receptor | 0.0 | molecule/$\mu m^2$ | | |
| 9 | LR*p | plasma membrane | Ligand-bound, phosphorylated active receptor | 0.0 | molecule/$\mu m^2$ | | |
| 10 | RpA | plasma membrane | Arrestin-bound, phosphorylated receptor | 0.0 | molecule/$\mu m^2$ | | |
| 11 | LRpA | plasma membrane | Ligand- and arrestin-bound, phosphorylated receptor | 0.0 | molecule/$\mu m^2$ | | |
| 12 | LR*pA | plasma membrane | Ligand- and arrestin-bound, phosphorylated active receptor | 0.0 | molecule/$\mu m^2$ | | |
| 13 | RipA | endosomes | Arrestin-bound, phosphorylated receptor | 0.0 | molecule/$\mu m^2$ | | |
| 14 | LRipA | endosomes | Ligand- and arrestin-bound, phosphorylated receptor | 0.0 | molecule/$\mu m^2$ | | |
| 15 | LRi*pA | endosomes | Ligand- and arrestin-bound, phosphorylated active receptor | 0.0 | molecule/$\mu m^2$ | | |
| 16 | Rip | endosomes | Phosphorylated receptor | 0.0 | molecule/$\mu m^2$ | | |
| 17 | LRip | endosomes | Ligand-bound, phosphorylated receptor | 0.0 | molecule/$\mu m^2$ | | |
| 18 | LRi*p | endosomes | Ligand-bound, phosphorylated active receptor | 0.0 | molecule/$\mu m^2$ | | |
| 19 | Ri | endosomes | Receptor | 0.0 | molecule/$\mu m^2$ | | |
| 20 | LRi | endosomes | Ligand-bound receptor | 0.0 | molecule/$\mu m^2$ | | |
| 21 | LRi* | endosomes | Ligand-bound, active receptor | 0.0 | molecule/$\mu m^2$ | | |
| 22 | G | plasma membrane | Heterotrimeric G protein | 40.0 | molecule/$\mu m^2$ | physiological expression | Ref. #41 |
| 23 | G_GTP | plasma membrane | GTP-bound α subunit of G protein | 0.0 | molecule/$\mu m^2$ | | |
| 24 | G_bg | plasma membrane | βγ subunit of G protein | 0.0 | molecule/$\mu m^2$ | | |
| 25 | G_GDP | plasma membrane | GDP-bound α subunit of G protein | 0.0 | molecule/$\mu m^2$ | | |
| 26 | RG | plasma membrane | G protein-bound receptor | 0.0 | molecule/$\mu m^2$ | | |
| 27 | LRG | plasma membrane | Ligand- and G protein-bound receptor | 0.0 | molecule/$\mu m^2$ | | |
| 28 | LR*G | plasma membrane | Ligand- and G protein-bound, active receptor | 0.0 | molecule/$\mu m^2$ | | |
| 29 | RG_bg | plasma membrane | βγ subunit-bound receptor | 0.0 | molecule/$\mu m^2$ | | |
| 30 | LRG_bg | plasma membrane | Ligand- and βγ subunit-bound receptor | 0.0 | molecule/$\mu m^2$ | | |
| 31 | LR*G_bg | plasma membrane | Ligand- and βγ subunit-bound, active receptor | 0.0 | molecule/$\mu m^2$ | | |
| 32 | PLC | plasma membrane | Phospholipase C | 10.0 | molecule/$\mu m^2$ | physiological expression | Ref. #41 |
| 33 | G_GTPPLC | plasma membrane | G_GTP–phospholipase C complex | 0.0 | molecule/$\mu m^2$ | | |
| 34 | G_GDPPLC | plasma membrane | G_GDP–phospholipase C complex | 0.0 | molecule/$\mu m^2$ | | |
| 35 | PI | plasma membrane | Phosphatidylinositol | 140000.0 | molecule/$\mu m^2$ | physiological concentration | Ref. #41 |
| 36 | PIP | plasma membrane | Phosphatidylinositol 4-phosphate | 3730.0 | molecule/$\mu m^2$ | physiological concentration, slightly changed to have equilibrium in the absence of ligand | Ref. #41 |
| 37 | PIP2 | plasma membrane | Phosphatidylinositol 4,5-bisphosphate | 5370.0 | molecule/$\mu m^2$ | physiological concentration, slightly changed to have equilibrium in the absence of ligand | Ref. #41 |
| 38 | boundPIP2 | plasma membrane | Protein-bound phosphatidylinositol 4,5-bisphosphate | 10740.0 | molecule/$\mu m^2$ | physiological concentration, slightly changed to have equilibrium in the absence of ligand | Ref. #41 |
| 39 | DAG | plasma membrane | Diacylglycerol | 20.0 | molecule/$\mu m^2$ | physiological concentration, slightly changed to have equilibrium in the absence of ligand | Ref. #41 |
| 40 | IP3 | plasma membrane | Inositol 1,4,5-trisphosphate | 12.5 | molecule/$\mu m^2$ | physiological concentration, slightly changed to have equilibrium in the absence of ligand | Ref. #41 |
| 41 | PKC | plasma membrane | Protein kinase C | 1000.0 | molecule/$\mu m^2$ | physiological concentration | Ref. #41 |
| 42 | PKCDAG | plasma membrane | DAG-bound protein kinase C | 0.0 | molecule/$\mu m^2$ | | |
| 43 | degradation | | Degradation of IP3 and DAG | 0.0 | molecule/$\mu m^2$ | | |

**Table S2. Reaction equations and rate constants of modeled reactions.** Table includes reaction index (from Fig. S16), equation and rate constant (with unit) for each reaction. Comments describe rationale for selecting rate constant in case of absence or discrepancy from literature reference.

| | Reaction | Compartment | Description | Value | Unit | Comment | Reference |
|---|---|---|---|---|---|---|---|
| **1** | L+R➔LR | plasma membrane | ligand association to receptor | 0.3 | 1/(μM*s) | Fast reaction | |
| **2** | LR➔L+R | plasma membrane | ligand dissociation from receptor | 0.0003 | 1/s | Was set to simulate a high-affinity ($K_D$=1nM) ligand | |
| **3** | LR➔LR* | plasma membrane | receptor activation; ligand-bound receptor | 0.3 | 1/s | Fast reaction | |
| **4** | LR*➔R+L | plasma membrane | ligand dissociation and receptor deactivation | 0.0003 | 1/s | Was set to simulate a high-affinity ($K_D$=1nM) ligand | |
| **5** | L+Rp➔LRp | plasma membrane | ligand association to phosphorylated receptor | 0.3 | 1/(μM*s) | Fast reaction | |
| **6** | LRp➔L+Rp | plasma membrane | ligand dissociation from phosphorylated receptor | 0.0003 | 1/s | Was set to simulate a high-affinity ($K_D$=1nM) ligand | |
| **7** | LRp➔LR*p | plasma membrane | receptor activation; ligand-bound, phosphorylated receptor | 0.3 | 1/s | Fast reaction | |
| **8** | LR*p➔Rp+L | plasma membrane | ligand dissociation and deactivation of phosphorylated receptor | 0.0003 | 1/s | Was set to simulate a high-affinity ($K_D$=1nM) ligand | |
| **9** | L+RpA➔LRpA | plasma membrane | ligand association to phosphorylated, arrestin-bound receptor | 0.3 | 1/(μM*s) | Fast reaction | |
| **10** | LRpA➔L+RpA | plasma membrane | ligand dissociation from phosphorylated, arrestin-bound receptor | 0.0003 | 1/s | Was set to simulate a high-affinity ($K_D$=1nM) ligand | |
| **11** | LRpA➔LR*pA | plasma membrane | receptor activation; ligand- and arrestin-bound, phosphorylated receptor | 0.3 | 1/s | Fast reaction | |
| **12** | LR*pA➔RpA+L | plasma membrane | ligand dissocation and deactivation of phosphorylated, arrestin-bound receptor | 0.0003 | 1/s | Was set to simulate a high-affinity ($K_D$=1nM) ligand | |
| **13** | Li+RipA➔LRipA | endosomes | ligand association to phosphorylated, arrestin-bound receptor | 0.3 | 1/(μM*s) | Fast reaction | |
| **14** | LRipA➔Li+RipA | endosomes | ligand dissociation from phosphorylated, arrestin-bound receptor | 0.0003 | 1/s | Was set to simulate a high-affinity ($K_D$=1nM) ligand | |
| **15** | LRipA➔LRi*pA | endosomes | receptor activation; ligand- and arrestin-bound, phosphorylated receptor | 0.3 | 1/s | Fast reaction | |
| **16** | LRi*pA➔Li+RipA | endosomes | ligand dissociation and deactivation of phosphorylated, arrestin-bound receptor | 0.0003 | 1/s | Was set to simulate a high-affinity ($K_D$=1nM) ligand | |
| **17** | Li+Rip➔LRip | endosomes | ligand association to phosphorylated receptor | 0.3 | 1/(μM*s) | Fast reaction | |
| **18** | LRip➔Li+Rip | endosomes | ligand dissociation from phosphorylated receptor | 0.0003 | 1/s | Was set to simulate a high-affinity ($K_D$=1nM) ligand | |
| **19** | LRip➔LRi*p | endosomes | receptor activation; phosphorylated receptor | 0.3 | 1/s | Fast reaction | |
| **20** | LRi*p➔Li+Rip | endosomes | ligand dissociation and deactivation of phosphorylated receptor | 0.0003 | 1/s | Was set to simulate a high-affinity ($K_D$=1nM) ligand | |
| **21** | Li+Ri➔LRi | endosomes | ligand association to receptor | 0.3 | 1/(μM*s) | Fast reaction | |
| **22** | LRi➔Li+Ri | endosomes | ligand dissociation from receptor | 0.0003 | 1/s | Was set to simulate a high-affinity ($K_D$=1nM) ligand | |
| **23** | LRi➔LRi* | endosomes | receptor activation; ligand-bound receptor | 0.3 | 1/s | Fast reaction | |
| **24** | LRi*➔Li+Ri | endosomes | ligand dissociation and receptor deactivation | 0.0003 | 1/s | Was set to simulate a high-affinity ($K_D$=1nM) ligand | |
| **25** | Rp➔R | plasma membrane | receptor dephosphorylation | 2.0 | 1/s | | Ref. #41 |
| **26** | R+PKCDAG➔Rp+PKCDAG | plasma membrane | receptor phosphorylation by PKC | 0.0 | 1/s | Not used in current simulations | |
| **27** | LRp➔LR | plasma membrane | receptor dephosphorylation; ligand-bound receptor | 2.0 | 1/s | | Ref. #41 |
| **28** | LR+PKCDAG➔LRp+PKCDAG | plasma membrane | receptor phosphorylation by PKC; ligand-bound receptor | 0.0 | 1/s | Not used in current simulations | |
| **29** | LR*p➔LR* | plasma membrane | receptor dephosphorylation; ligand-bound, active receptor | 2.0 | 1/s | | Ref. #41 |
| **30** | LR*➔LR*p | plasma membrane | receptor phosphorylation by GRK; ligand-bound, active receptor | 2.0 | 1/s | | was set between values of Ref. #41 and #42 |
| **31** | LR*+PKCDAG➔LR*p+PKCDAG | plasma membrane | receptor phosphorylation by PKC; ligand-bound, active receptor | 0.0 | 1/s | Not used in current simulations | |
| **32** | RpA➔Rp+A | plasma membrane | arrestin dissociation from phosphorylated receptor | 0.01 | 1/s | 33.3× faster than reaction 34 | |
| **33** | LRpA➔LRp+A | plasma membrane | arrestin dissociation from phosphorylated, ligand-bound receptor | 0.01 | 1/s | 33.3× faster than reaction 34 | |
| **34** | LR*pA➔LR*p+A | plasma membrane | arrestin dissociation from phosphorylated, ligand-bound, active receptor | 0.0003 | 1/s | | was set between k13 and k14 in Ref. #42 |
| **35** | LR*p+A➔LR*pA | plasma membrane | arrestin association to phosphorylated, ligand-bound, active receptor | 0.000001 | $\mu m^2$/s | | based on k12 in Ref. #42 and arrestin concentrations, slightly modified |
| **36** | LRp+A➔LRpA | plasma membrane | arrestin association to phosphorylated, ligand-bound receptor | 0.0000003 | $\mu m^2$/s | 3.33× slower than reaction 35 | |
| **37** | Rp+A➔RpA | plasma membrane | arrestin association to phosphorylated receptor | 0.0000003 | $\mu m^2$/s | 3.33× slower than reaction 35 | |
| **38** | LR*pA➔LRi*pA | | internalization of phosphorylated, ligand- and arrestin-bound active receptor | 0.0073 | 1/s | | Ref. #28 |
| **39** | LRpA➔LRipA | | internalization of phosphorylated, ligand- and arrestin-bound receptor | 0.0073 | 1/s | | Ref. #28 |
| **40** | RpA➔RipA | | internalization of phosphorylated, arrestin-bound receptor | 0.0073 | 1/s | | Ref. #28 |
| **41** | RipA➔Rip+A | endosomes | arrestin dissociation from phosphorylated receptor | 0.01 | 1/s | 33.3× faster than reaction 43 | |
| **42** | LRipA➔LRip+A | endosomes | arrestin dissociation from phosphorylated, ligand-bound receptor | 0.01 | 1/s | 33.3× faster than reaction 43 | |
| **43** | LRi*pA➔LRi*p+A | endosomes | arrestin dissociation from phosphorylated, ligand-bound, active receptor | 0.0003 | 1/s | | was set between k13 and k14 in Ref. #42 |
| **44** | LRi*p+A➔LRi*pA | endosomes | arrestin association to phosphorylated, ligand-bound, active receptor | 0.000001 | $\mu m^2$/s | | based on k12 in Ref. #42 and arrestin concentrations, slightly modified |
| **45** | LRip+A➔LRipA | endosomes | arrestin association to phosphorylated, ligand-bound receptor | 0.0000003 | $\mu m^2$/s | 3.33× slower than reaction 44 | |
| **46** | Rip+A➔RipA | endosomes | arrestin association to phosphorylated receptor | 0.0000003 | $\mu m^2$/s | 3.33× slower than reaction 44 | |
| **47** | Rip➔Ri | endosomes | receptor dephosphorylation | 2.0 | 1/s | | Ref. #41 |
| **48** | LRip➔LRi | endosomes | receptor dephosphorylation; ligand-bound receptor | 2.0 | 1/s | | Ref. #41 |
| **49** | LRi*p➔LRi* | endosomes | receptor dephosphorylation; ligand-bound, active receptor | 2.0 | 1/s | | Ref. #41 |
| **50** | LRi*➔LRi*p | endosomes | receptor phosphorylation by GRK; ligand-bound, active receptor | 0.02 | 1/s | 100× slower than reaction 30 | |
| **51** | L+RG➔LRG | plasma membrane | ligand association to heterotrimeric G protein-bound receptor | 0.3 | 1/(μM*s) | Fast reaction | |
| **52** | LRG➔L+RG | plasma membrane | ligand dissociation from heterotrimeric G protein-bound receptor | 0.0003 | 1/s | Was set to simulate a high-affinity ($K_D$=1nM) ligand | |
| **53** | LRG➔LR*G | plasma membrane | receptor activation; heterotrimeric G protein-bound receptor | 0.3 | 1/s | Fast reaction | |
| **54** | LR*G➔L+RG | plasma membrane | ligand dissociation and deactivation of heterotrimeric G protein-bound receptor | 0.0003 | 1/s | Was set to simulate a high-affinity ($K_D$=1nM) ligand | |
| **55** | L+RG_bg➔LRG_bg | plasma membrane | ligand association to βγ-subunit-bound receptor | 0.3 | 1/(μM*s) | Fast reaction | |
| **56** | LRG_bg➔L+RG_bg | plasma membrane | ligand dissociation from βγ-subunit-bound receptor | 0.0003 | 1/s | Was set to simulate a high-affinity ($K_D$=1nM) ligand | |

| | Reaction | Compartment | Description | Value | Unit | Comment | |
|---|---|---|---|---|---|---|---|
| **57** | LRG_bg→LR*G_bg | plasma membrane | receptor activation; βγ-subunit-bound receptor | 0.3 | 1/s | Fast reaction | |
| **58** | LR*G_bg→L+RG_bg | plasma membrane | ligand dissociation and deactivation of βγ-subunit-bound receptor | 0.0003 | 1/s | Was set to simulate a high-affinity ($K_D$=1nM) ligand | |
| **59** | RG→R+G | plasma membrane | dissociation of heterotrimeric G protein from receptor | 6.8 | 1/s | | Ref. #43 |
| **60** | R+G→RG | plasma membrane | heterotrimeric G protein association to receptor | 0.00027 | $\mu m^2$/s | | Ref. #43 |
| **61** | LRG→LR+G | plasma membrane | dissociation of heterotrimeric G protein from ligand-bound receptor | 6.8 | 1/s | | Ref. #43 |
| **62** | LR+G→LRG | plasma membrane | heterotrimeric G protein association to ligand-bound receptor | 0.00027 | $\mu m^2$/s | | Ref. #43 |
| **63** | LR*G→LR*+G | plasma membrane | dissociation of heterotrimeric G protein from ligand-bound, active receptor | 0.68 | 1/s | | Ref. #43 |
| **64** | LR*+G→LR*G | plasma membrane | heterotrimeric G protein association to ligand-bound, active receptor | 0.0027 | $\mu m^2$/s | | Ref. #43 |
| **65** | RG_bg+G_GDP→RG | plasma membrane | reassociation of heterotrimeric G protein complex on receptor | 1.0 | $\mu m^2$/s | | Ref. #43 |
| **66** | RG→RG_bg+G_GTP | plasma membrane | activation of heterotrimeric G protein complex by receptor | 0.000015 | 1/s | | Ref. #43 |
| **67** | LRG_bg+G_GDP→LRG | plasma membrane | reassociation of heterotrimeric G protein complex on ligand-bound receptor | 1.0 | $\mu m^2$/s | | Ref. #43 |
| **68** | LRG→LRG_bg+G_GTP | plasma membrane | activation of heterotrimeric G protein complex by ligand-bound receptor | 0.000015 | 1/s | | Ref. #43 |
| **69** | LR*G_bg+G_GTP→LR*G | plasma membrane | reassociation of heterotrimeric G protein complex on ligand-bound, active receptor | 1.0 | 1/(μM*s) | | Ref. #43 |
| **70** | LR*G→LR*G_bg+G_GTP | plasma membrane | activation of heterotrimeric G protein complex by ligand-bound, active receptor | 0.65 | 1/s | | Ref. #43 |
| **71** | R+G_bg→RG_bg | plasma membrane | association of βγ-subunit to receptor | 0.00027 | 1/(μM*s) | | Ref. #43 |
| **72** | RG_bg→R+G_bg | plasma membrane | dissociation of βγ-subunit from receptor | 6.8 | 1/s | | Ref. #43 |
| **73** | LR+G_bg→LRG_bg | plasma membrane | association of βγ-subunit to ligand-bound receptor | 0.00027 | $\mu m^2$/s | | Ref. #43 |
| **74** | LRG_bg→LR+G_bg | plasma membrane | dissociation of βγ-subunit from ligand-bound receptor | 6.8 | 1/s | | Ref. #43 |
| **75** | LR*+G_bg→LR*G_bg | plasma membrane | association of βγ-subunit to ligand-bound, active receptor | 0.0027 | $\mu m^2$/s | | Ref. #43 |
| **76** | LR*G_bg→LR*+G_bg | plasma membrane | dissociation of βγ-subunit from ligand-bound, active receptor | 0.68 | 1/s | | Ref. #43 |
| **77** | G_GTP+PLC→G_GTPPLC | plasma membrane | association of GTP-bound Gα subunit to PLC | 1.0 | $\mu m^2$/s | | Ref. #43 |
| **78** | G_GDPPLC→G_GDP+PLC | plasma membrane | dissocation of GDP-bound Gα subunit from PLC | 0.71 | 1/s | | Ref. #43 |
| **79** | G_GTP→G_GDP | plasma membrane | GTPase activity of Gα subunit | 0.026 | 1/s | | Ref. #43 |
| **80** | G_GDPPLC→G_GTPPLC | plasma membrane | GDP–GTP exchange of PLC-Gα subunit complex | 4.7 | 1/s | | Ref. #43 |
| **81** | G_GTPPLC→G_GDPPLC | plasma membrane | GTPase activity of Gα subunit in PLC-Gα subunit complex | 15.0 | 1/s | | Ref. #43 |
| **82** | G_GDP+G_bg→G | plasma membrane | reassociation of heterotrimeric G protein complex | 1.0 | $\mu m^2$/s | | Ref. #43 |
| **83** | G→G_GTP+G_bg | plasma membrane | spontaneous activation of heterotrimeric G protein complex | 0.000015 | 1/s | | Ref. #43 |
| **84** | PI→PIP | plasma membrane | Phosphatidylinositol phosphorylation | 0.00078 | 1/s | | Ref. #44 |
| **85** | PI+PKCDAG→PIP+PKCDAG | plasma membrane | PKC-dependent phosphatidylinositol phosphorylation | 0.0000039 | 1/s | | Ref. #44 |
| **86** | PIP→PI | plasma membrane | Phosphatidylinositol 4-phosphate dephosphorylation | 0.03 | 1/s | | Ref. #44 |
| **87** | PIP→PIP2 | plasma membrane | Phosphatidylinositol 4-phosphate phosphorylation | 0.06 | 1/s | | Ref. #44 |
| **88** | PIP+PKCDAG→PIP2+PKCDAG | plasma membrane | PKC-dependent phosphatidylinositol 4-phosphate phosphorylation | 0.00012 | 1/s | | Ref. #44 |
| **89** | PIP2→PIP | plasma membrane | Phosphatidylinositol 4,5-bisphosphate dephosphorylation | 0.042 | 1/s | | Ref. #44 |
| **90** | PIP2→boundPIP2 | plasma membrane | Phosphatidylinositol 4,5-bisphosphate association to proteins | 2.0 | 1/s | | Ref. #44 |
| **91** | boundPIP2→PIP2 | plasma membrane | Phosphatidylinositol 4,5-bisphosphate dissociation from proteins | 1.0 | 1/s | | Ref. #44 |
| **92** | PIP2+G_GTPPLC→DAG+IP3+G_GTPPLC | plasma membrane | Phosphatidylinositol 4,5-bisphosphate cleavage by PLC | 0.6 | 1/s | | Ref. #44 |
| **93** | PKCDAG→PKC+DAG | plasma membrane | Dissociation of PKC from DAG | 0.06 | 1/s | | Ref. #44 |
| **94** | DAG+PKC→PKCDAG | plasma membrane | PKC association to DAG | 0.00002 | $\mu m^2$/s | | Ref. #44 |
| **95** | DAG→degradation | | DAG degradation | 0.05 | 1/s | | Ref. #44 |
| **96** | IP3→degradation | | IP3 degradation | 0.08 | 1/s | | Ref. #44 |

#### Table S3. Applied plasmid DNA amounts in the distinct BRET setups

| BRET setup | Plasmid #1 (amount in µg/well) | Plasmid #2 (amount in µg/well) | Plasmid #3 (amount in µg/well) | Plasmid #4 (amount in µg/well) | Transfection method |
|---|---|---|---|---|---|
| AT1R - β-arrestin1 interaction | AT1R–Rluc (0.075) | β-arrestin1–Venus (0.075) | pcDNA3.1 (0.15) | - | Lipofectamine 2000 |
| AT1R - β-arrestin2 interaction | AT1R–Rluc (0.075) | β-arrestin2–Venus (0.075) | pcDNA3.1 (0.15) | - | Lipofectamine 2000 |
| AT1R - β-arrestin2 interaction, +Dyn-K44A | AT1R–Rluc (0.075) | β-arrestin2–Venus (0.075) | HA–dynamin2A-K44A (0.15) | - | Lipofectamine 2000 |
| AT1R - β-arrestin2 interaction, +PtdIns(4,5)P2 depletion system | AT1R–Rluc (0.05) | β-arrestin2–Venus (0.05) | L10–FRB–T2A–FKBP–5ptase (0.2) | - | Lipofectamine 2000 |
| Gi1 activation (TRUPATH) | AT1R (0.075) | Gαi1–Rluc8 (0.075) | Gβ3 (0.075) | Gγ9–GFP2 (0.075) | Lipofectamine 2000 |
| Gi2 activation (TRUPATH) | AT1R (0.075) | Gαi2–Rluc8 (0.075) | Gβ3 (0.075) | Gγ8–GFP2 (0.075) | Lipofectamine 2000 |
| Gi3 activation (TRUPATH) | AT1R (0.075) | Gαi3–Rluc8 (0.075) | Gβ3 (0.075) | Gγ9–GFP2 (0.075) | Lipofectamine 2000 |
| GoA activation (TRUPATH) | AT1R (0.075) | GαoA–Rluc8 (0.075) | Gβ3 (0.075) | Gγ8–GFP2 (0.075) | Lipofectamine 2000 |
| GoB activation (TRUPATH) | AT1R (0.075) | GαoB–Rluc8 (0.075) | Gβ3 (0.075) | Gγ8–GFP2 (0.075) | Lipofectamine 2000 |
| Gs activation (TRUPATH) | AT1R (0.075) | GαsL–Rluc8 (0.075) | Gβ1 (0.075) | Gγ1–GFP2 (0.075) | Lipofectamine 2000 |
| Gq activation (TRUPATH) | AT1R (0.075) | Gαq–Rluc8 (0.075) | Gβ3 (0.075) | Gγ9–GFP2 (0.075) | Lipofectamine 2000 |
| G11 activation (TRUPATH) | AT1R (0.075) | Gα11–Rluc8 (0.075) | Gβ3 (0.075) | Gγ13–GFP2 (0.075) | Lipofectamine 2000 |
| G12 activation (TRUPATH) | AT1R (0.075) | Gα12–Rluc8 (0.075) | Gβ3 (0.075) | Gγ9–GFP2 (0.075) | Lipofectamine 2000 |
| G13 activation (TRUPATH) | AT1R (0.075) | Gα13–Rluc8 (0.075) | Gβ3 (0.075) | Gγ9–GFP2 (0.075) | Lipofectamine 2000 |
| PM PtdIns(4,5)P2 level | AT1R (0.075) | L10–Venus–T2A–PLCδ1PH–SLuc (0.075) | pcDNA3.1 (0.15) | - | Lipofectamine 2000 |
| PM PtdIns(4,5)P2 level, +Dyn-K44A | AT1R (0.075) | L10–Venus–T2A–PLCδ1PH–SLuc (0.075) | HA–dynamin2A-K44A (0.15) | - | Lipofectamine 2000 |
| PM PtdIns(4,5)P2 level, +β-arrestin2 | AT1R (0.075) | L10–Venus–T2A–PLCδ1PH–SLuc (0.075) | β-arrestin2 (0.15) | - | Lipofectamine 2000 |
| AT1R activation-induced β-arrestin2 recruitment to the PM | AT1R (0.1) | PM–NanoLuc (0.001) | β-arrestin2–Venus (0.1) | pcDNA3.1 (0.15) | Lipofectamine 2000 |
| AT1R activation-induced β-arrestin2 recruitment to the EEs | AT1R (0.1) | NanoLuc–EE (0.001) | β-arrestin2–Venus (0.1) | pcDNA3.1 (0.15) | Lipofectamine 2000 |
| β2AR - β-arrestin2 interaction | β2AR–SLuc (0.025) | β-arrestin2–Venus (0.075) | pcDNA3.1 (0.15) | GRK2 (0.05) | Lipofectamine 2000 |
| β2AR - β-arrestin2 interaction, +Dyn-K44A | β2AR–SLuc (0.025) | β-arrestin2–Venus (0.075) | HA–dynamin2A-K44A (0.15) | GRK2 (0.05) | Lipofectamine 2000 |
| β2AR-3S - β-arrestin2 interaction | β2AR-3S–SLuc (0.025) | β-arrestin2–Venus (0.075) | pcDNA3.1 (0.15) | GRK2 (0.05) | Lipofectamine 2000 |
| β2AR-3S - β-arrestin2 interaction, +Dyn-K44A | β2AR-3S–SLuc (0.025) | β-arrestin2–Venus (0.075) | HA–dynamin2A-K44A (0.15) | GRK2 (0.05) | Lipofectamine 2000 |
| AT1R internalization (disappearence from the PM) | AT1R–Rluc (0.075) | L10–Venus (0.075) | pcDNA3.1 (0.15) | - | Lipofectamine 2000 |
| AT1R internalization (disappearence from the PM), +β-arrestin2 | AT1R–Rluc (0.075) | L10–Venus (0.075) | β-arrestin2 (0.15) | - | Lipofectamine 2000 |
| AT1R internalization (appearence in the EEs) | AT1R–Rluc (0.075) | Venus–Rab5 (0.075) | pcDNA3.1 (0.15) | - | Lipofectamine 2000 |
| AT1R internalization (appearence in the EEs), +β-arrestin2 | AT1R–Rluc (0.075) | Venus–Rab5 (0.075) | β-arrestin2 (0.15) | - | Lipofectamine 2000 |
| AT1R–β-arrestin2–MEK1 complex formation | AT1R–Rluc8 (0.01) | Venus–MEK1–FLAG (0.1) | β-arrestin2 (0.1) | - | Lipofectamine 2000 |
| AT1R–β-arrestin2–ERK2 complex formation | AT1R–Rluc8 (0.01) | FLAG–ERK2–Venus (0.1) | β-arrestin2 (0.1) | - | Lipofectamine 2000 |
| Competitive kinetic ligand binding measurements | AT1R (0.15) | GLuc–PM (0.01) | β-arrestin2 (0.06) | HA–dynamin2A-K44A (0.04) | Calcium phosphate |

**Table S4. Table of applied luciferase substrates and filters in BRET measurements**

| **Donor** | **Substrate** | **Acceptor** | **Filter1** | **Filter2** |
|---|---|---|---|---|
| *Renilla* luciferase (humanized, SLuc, RLuc8) | coelenterazine *h* (5 μM) | Venus fluorescent protein | 480/20 nm | 530/20 nm |
| *Renilla* luciferase (RLuc8) | Prolume Purple (1 μM) | GFP2 fluorescent protein | 425/50 nm | 515/30 nm |
| Nanoluciferase (NanoLuc) | coelenterazine *h* (5 μM) | Venus fluorescent protein | 460/20 nm | 530/20 nm |
| *Gaussia* luciferase (GLucM23) | native coelenterazine (5 μM) | TAMRA fluorescent dye | 480/20 nm | 610/60 nm |
